# Apical PAR complex proteins protect against epithelial assaults to create a continuous and functional intestinal lumen

**DOI:** 10.1101/2020.10.28.359299

**Authors:** Maria D. Sallee, Melissa A. Pickett, Jessica L. Feldman

## Abstract

Sustained polarity and adhesion of epithelial cells is essential for the protection of our organs and bodies, and this epithelial integrity emerges during organ development amidst numerous morphogenetic assaults. Using the developing *C. elegans* intestine as an *in vivo* model, we investigated how epithelial cells maintain integrity through cell division and elongation to build a functional tube. Live-imaging revealed that apical PAR complex proteins PAR-6/Par6 and PKC-3/aPkc remained apical during mitosis while apical microtubules and microtubule-organizing center (MTOC) proteins were transiently removed. Intestine-specific depletion of PAR-6, PKC-3, and the aPkc regulator CDC-42/Cdc42 caused persistent gaps in the apical MTOC as well as in other apical and junctional proteins after cell division and in non-dividing cells that elongated. Upon hatching, gaps coincided with luminal constrictions that blocked food, and larvae arrested and died. Thus, the apical PAR complex maintains apical and junctional continuity to construct a functional intestinal tube.

## INTRODUCTION

Epithelia are dynamic tissues composed of highly polarized and adherent epithelial cells that line organs and act as selective barriers and transporters, provide mechanical resiliency to organs, and separate the inside of animal bodies from the outside world. To form and function correctly, epithelia must maintain their integrity during development and homeostasis as they are challenged by various assaults including mitosis, cell shape change, intercalation, cell death, and mechanical forces. How epithelial primordia maintain their integrity *in vivo* in the face of these assaults to build a functional organ is not well understood.

Apicobasal polarity is a hallmark of epithelia, and tissue integrity requires that polarity and adhesion be coordinated throughout the tissue (Pickett et al., 2019); outward facing apical surfaces must be aligned and junctional complexes (“junctions”) must form between neighboring cells. Apical surfaces frequently anchor microtubules (Sanchez and Feldman, 2017) and have important functional specializations, such as microvilli and cilia, while basolateral domains are important for cell-cell contact and adhesion to overlying basement membranes. Junctions form belt-like bands at the cell periphery separating the apical and basolateral domains. Adherens junctions provide adhesion, anchor the actin cytoskeleton, and impart mechanical strength by connecting the cytoskeletons of adjacent cells; tight/septate junctions form a diffusion barrier through extracellular binding of junctional proteins between neighboring cells (Rusu and Georgiou, 2020), contributing to epithelial integrity. The establishment and maintenance of apical and basolateral domains as well as the formation and maturation of junctions depends on the apical PAR complex proteins PAR-3/Par3, PAR-6/Par6, PKC-3/aPkc, and CDC-42/Cdc42 (hereafter referred to as the ‘PAR complex’ for simplicity) in many tissues and organisms (Pickett et al., 2019). The scaffold Par3 recruits another scaffold Par6, which binds to and localizes interdependently with the kinase aPkc to the apical surface, and the Rho-family GTPase Cdc42 can displace Par3 and activate aPkc kinase activity (Achilleos et al., 2010; Totong et al., 2007)(Harris and Peifer, 2005; Hutterer et al., 2004; Joberty et al., 2000; Kim et al., 2009; Lin et al., 2000)(Rodriguez et al., 2017). Active aPkc phosphorylates and removes basolateral proteins Lgl and Par1 from the apical domain, and mutual inhibition between apical and basolateral proteins maintains and refines the apical and basolateral domains (Pickett et al., 2019).

The PAR complex also promotes apical junction formation and maintenance (Georgiou et al., 2008; Montoyo-Rosario et al., 2020; Totong et al., 2007; Wallace et al., 2010; Zilberman et al., 2017). Junctions form at cell-cell contacts first as immature E-cadherin clusters that recruit additional proteins. Junctions then mature by condensing into distinct continuous bands of E-cadherin-based adherens junctions and occludin/claudin-based tight junctions (Rusu and Georgiou, 2020). PAR complex proteins promote tight junction assembly and regulate E-cadherin endocytosis, which is essential for adherens junction plasticity and maintenance, and Cdc42 regulates the actin cytoskeleton which stabilizes junctions (Mack and Georgiou, 2014).

Following polarization and junction establishment, epithelial tissues remain dynamic during development and homeostasis as cells can divide, die, or change their shape, size, or arrangement. Such assaults on their integrity require epithelial cells to remodel existing polarized structures or build new ones. In most epithelia, cell division physically cleaves their polarized domains and daughters must build junctions at the new cell-cell contact while maintaining correct apicobasal polarity. Division also requires rearrangement of the microtubule cytoskeleton. Parallel arrays of microtubules emanate from the apical microtubule-organizing center (MTOC) in interphase epithelial cells, where they direct intracellular vesicle transport and maintain apicobasal polarity (Sanchez and Feldman, 2017). During mitosis, cells temporarily rearrange their microtubules into radial arrays around the centrosomal MTOC and then return them to the apical surface (Pease and Tirnauer, 2011; Yang and Feldman, 2015). Epithelial cells also change their shape and size during development and homeostasis, expanding or shrinking their apical surfaces in concert with their junctions to maintain their integrity. Apical constriction shrinks the apical surface and generates epithelial folds that drive *Drosophila* gastrulation and vertebrate neurulation (Martin and Goldstein, 2014). Cells can also undergo apical emergence, extending to join the apical side of the epithelium and expanding their apical domains and junctions, as seen in multiciliated cells of the *Xenopus* epidermis (Sedzinski et al., 2016). Changes to the apical surface generally require the addition or removal of apical membrane through vesicle trafficking and force applied by actin and myosin on junctions (Martin and Goldstein, 2014). Importantly, PAR complex proteins play critical roles in spindle orientation in dividing cells (Mack and Georgiou, 2014) and guide microtubule localization to the apical surface during polarity establishment (Feldman and Priess, 2012), and several studies have found requirements for PAR complex proteins in regulating actin dynamics during apical constriction (David et al., 2010; Ishiuchi and Takeichi, 2011; Walck-Shannon et al., 2016; Zilberman et al., 2017). However, the role of PAR complex proteins in restoring apical microtubule organization and in maintaining the apical domain and junctions during morphogenesis is less well understood due to their early roles in zygotes and in embryonic epithelia (Achilleos et al., 2010; Gotta et al., 2001; Harris and Peifer, 2005; Kay and Hunter, 2001; Montoyo-Rosario et al., 2020; Plusa et al., 2005; Tabuse et al., 1998; Totong et al., 2007; Watts et al., 1996; Zilberman et al., 2017).

Recent advances in tissue-specific protein depletion allow robust spatial and temporal control of protein removal, enabling us to study the later roles of polarity proteins in specific tissues (Armenti et al., 2014; Sallee et al., 2018; Wang et al., 2017; Zhang et al., 2015). The developing *C. elegans* intestine provides a genetically accessible *in vivo* context to study how an epithelial primordium maintains tissue integrity through developmental assaults. The sixteen cells of the intestinal primordium polarize and establish their apical domains around a central midline, the future site of the intestinal lumen. Two anterior and two posterior cells undergo a final embryonic division, and all twenty resulting cells must actively remodel and expand their apical surfaces and junctions as the intestine elongates in order to maintain apical and junctional continuity along the midline. By the time embryos hatch, the intestine is a tube with a continuous open lumen and intact barrier function, capable of digestion. We can thus use the *C. elegans* intestine to investigate how epithelial integrity is maintained through cell division and elongation, and the consequence of disrupting that integrity on organ morphology and function and organismal health.

Here, we found that PAR complex proteins are required during intestinal development to build a functional intestine with a continuous open lumen. Intestine-specific depletion of PAR-6, PKC-3, and CDC-42 resulted in gaps along the midline in apical MTOC proteins, both where cells had divided and where cells were elongating. In PAR-6-depleted intestines, gaps in MTOC proteins correlated with gaps in apical and junctional proteins, and basolateral proteins failed to be excluded from the midline. The discontinuities in apical and junctional proteins became more numerous as elongation progressed, and larvae arrested upon hatching with functionally obstructed intestines showing gaps in the lumen, apical surfaces, and junctions. Together, these results demonstrate the importance of these highly conserved PAR complex proteins during development in remodeling the apical domain and junctions to keep continuity as intestinal cells divide and expand their apical surfaces.

## RESULTS

### Cells within the polarized C. elegans intestine elongate and divide

Following polarization, cells in the developing *C. elegans* intestine divide, intercalate, and elongate during morphogenesis. Through division and elongation, intestinal cells maintain apical and junctional continuity across neighboring cells, and thus serve as a good *in vivo* model to understand the mechanisms epithelial cells use to buffer against these assaults. The sixteen-cell primordial intestine (E16) is arranged into two tiers of ten dorsal and six ventral cells. These cells begin to polarize soon after they are born, building a continuous apical domain along a central midline, the future lumen (Leung et al., 1999). Adherens and septate junctions form a single ‘apical junction’ that localizes around the apical domain between left/right, dorsal/ventral, and anterior/posterior pairs of cells in a ladder-like arrangement along the midline ((Costa et al., 1998; Labouesse, 2006; Leung et al., 1999), Supp. Fig. 1). Two pairs of anterior and posterior cells, referred to here as “star cells,” undergo an additional round of mitosis after polarity is established while the other twelve “non-star cells” remain in interphase. The intestine more than doubles its midline length from a polarized primordium at bean stage to the 1.5- to 1.8-fold stage (Fig. 1A-C). Elongation continues and the lumen is formed at the midline. Embryos hatch as L1 (first larval stage) larvae with a functional intestine (Fig. 1C), built from a series of nine stacked “int” rings with a continuous central hollow lumen through which food passes (Supp. Fig. 1). The apical surfaces of all the cells face the lumen where they secrete digestive enzymes and absorb nutrients through microvilli, and junctional complexes maintain cell adhesion and barrier function while allowing the open lumen to run unobstructed between the intestinal rings (Dimov and Maduro, 2019).

**Figure 1.**
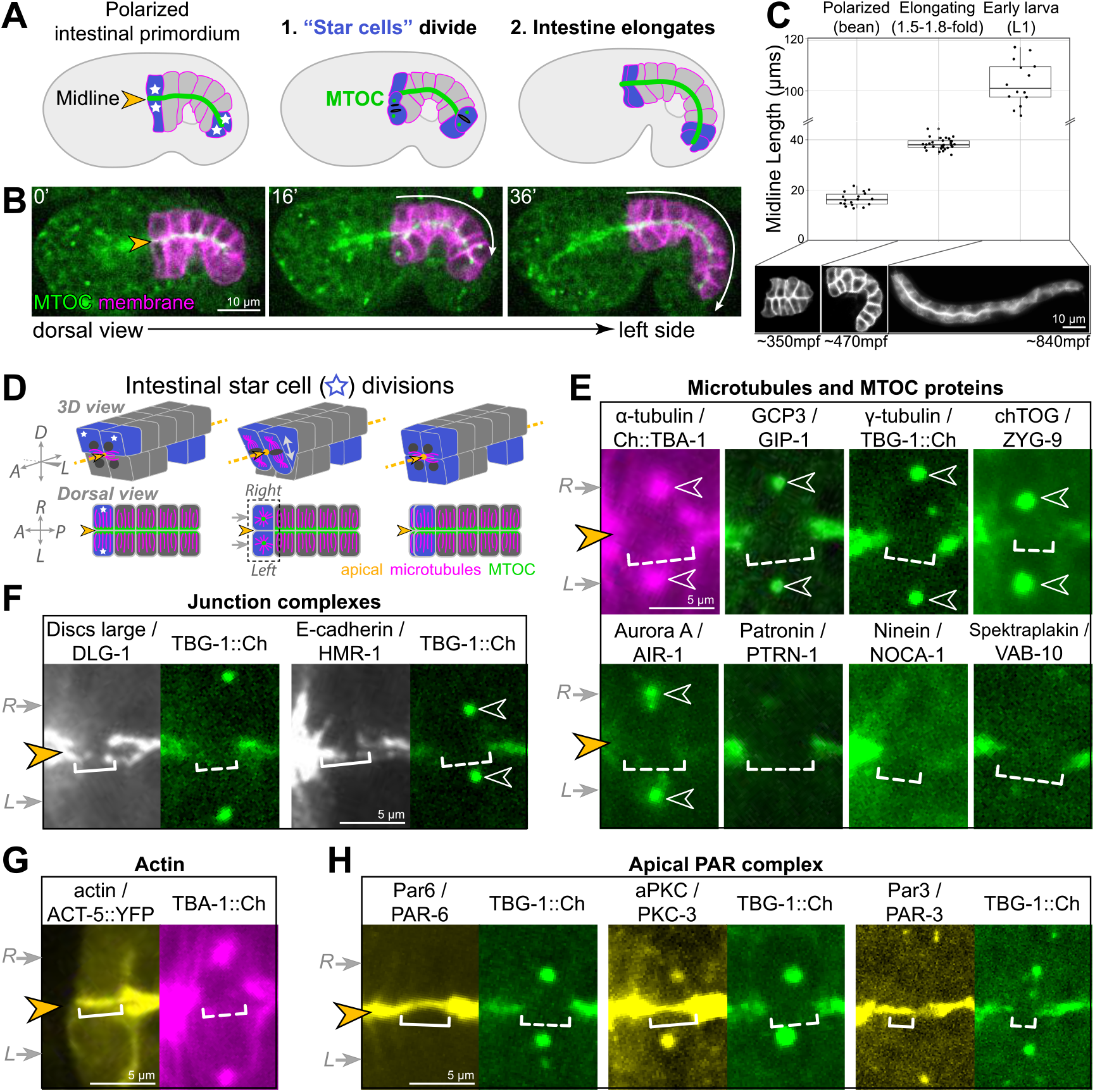
Cell division and elongation challenge the epithelial integrity of the developing C. elegans intestine. (A, B) A cartoon schematic (A) and corresponding live time course (B) of an embryo expressing an apical MTOC marker TBG-1::mCherry (green, orange arrowhead) and a membrane marker intestinal GFP::CAAX (magenta); the anterior and posterior “star cells” divide (blue) and the apical surface elongates (white arrow) as the polarized intestinal primordium develops into an intestine. (C) Top: Graph of intestinal apical length in newly polarized primordia (bean stage, average length = 16.5±2.6 μm, n = 18), ~1.5-2 hours after the start of intestinal elongation (1.5-fold to 1.8-fold, average length = 38.2±2.2 μm, n = 30), and upon hatching in the L1 larval stage (average length = 105.0±11.1 μm, n = 15). Below: Corresponding intestinal GFP::CAAX images. (D) Cartoon schematic of the anterior star cell divisions illustrating their dorsoventral division in 3D (top) and from a dorsal view (bottom). Dotted line indicates viewing angle of images in panels E-H. (E-H) Live imaging of indicated proteins relative to TBG-1/γ-tubulin::mCherry (green) or mCherry::TBA-1/α-tubulin (magenta) and the midline (orange arrowhead) when the right and left cells divided synchronously (gray arrows). Note that in some cases a midline gap (dashed bracket) formed, and in other cases no gap formed (solid bracket). All images are maximum intensity Z-projections that capture centrosomes and/or the intestinal midline. Scale bar = 10 μm in B, C. Scale bars = 5 μm in E-H.

### PAR complex proteins remain apical during mitosis, when the apical MTOC is transiently removed

We first examined the localization of different polarized proteins during mitosis to determine the impact of cell division on their localization. Apically organized microtubules are one hallmark of epithelial cells that is necessarily affected by mitosis. Apical microtubules and the microtubule nucleator γ-Tubulin Ring Complex (γ-TuRC) transiently leave the apical surface when star cells divide and centrosomes are reactivated as MTOCs to build the mitotic spindle (Pease and Tirnauer, 2011; Yang and Feldman, 2015). The loss of apical microtubules is easiest to see when star cells divide at the same time. Both apical surfaces lose apical microtubules during mitosis, creating a visible “midline gap” in apical MTOC proteins as both the left and right cells reactivate centrosomal MTOCs; the midline gap is then refilled upon mitotic exit (Fig. 1D, 1E; (Yang and Feldman, 2015)). We found that in addition to microtubules and the γ-TuRC components TBG-1/γ-tubulin and GIP-1/GCP3, all MTOC and microtubule-associated proteins we examined also left the apical surface during mitosis (Fig. 1E). Some MTOC proteins localized to the active centrosomes (γ-TuRC, ZYG-9/chTOG, AIR-1/Aurora A), while other proteins did not and likely became cytoplasmic (NOCA-1/Ninein, PTRN-1/CAMSAP, VAB-10B/Spectraplakin). The observed removal of apical MTOC proteins during mitosis suggests the complete inactivation of the apical surface as an MTOC. Junctional proteins retained some localization during mitosis, albeit reduced as compared with interphase neighboring cells.

Localization of the septate-like junction-associated protein DLG-1/Discs large was most strongly reduced during mitosis, and its intensity was frequently undetectable during the star cell divisions (Fig. 1F). The adherens junction protein HMR-1/E-cadherin remained localized, though its intensity was reduced compared to the non-dividing “non-star” intestinal cells, appearing as thin fluorescent threads as new cell boundaries formed. The other apical proteins examined, ACT-5/actin and three members of the PAR complex PAR-3/Par3, PAR-6/Par6, and PKC-3/aPkc, all remained apically localized during mitosis (Fig. 1G, H). Therefore, we hypothesized that PAR complex proteins actively mark the apical surface during mitotic remodeling to return apical MTOC function after division, thereby restoring a continuous apical MTOC.

### PAR-6 and PKC-3 are required for the return of apical MTOC function after mitosis

To test our hypothesis that the PAR complex acts as an apical memory mark, we asked if removing PAR-6 or its well-characterized binding partner PKC-3 during the star cell divisions would disrupt the return of the MTOC to the apical surface. To avoid the embryonic lethality caused by loss of PAR-6 or PKC-3 in the zygote or in the skin (Montoyo-Rosario et al., 2020; Tabuse et al., 1998; Totong et al., 2007; Watts et al., 1996), we used the ZIF-1/ZF degradation system to deplete each protein only in the intestine (“gut(−)”)(Armenti et al., 2014; Sallee et al., 2018). Briefly, we used CRISPR/Cas9 to add a ZF::GFP tag to endogenous PKC-3 and PAR-6; the ZF1 degron (“ZF”) facilitated efficient degradation by the E3 ligase component ZIF-1, and GFP allowed visualization of protein localization and depletion. We used intestine-specific promoters to express ZIF-1 prior to polarity establishment (*elt-2*p in the 4-intestinal cell “E4” stage and *ifb-2*p in the eight-intestinal cell “E8” stage), ensuring robust depletion only in the gut well before star cell divisions and intestinal elongation (Supp. Fig. 2). Unlike PAR-3, PAR-6 and PKC-3 are not required to establish apicobasal polarity in the intestine ((Achilleos et al., 2010; Totong et al., 2007), Supp. Fig. 2), so we could degrade PAR-6 in early stages of intestinal development without disrupting polarity establishment itself and ensure robust depletion prior to the star cell divisions.

**Figure 2.**
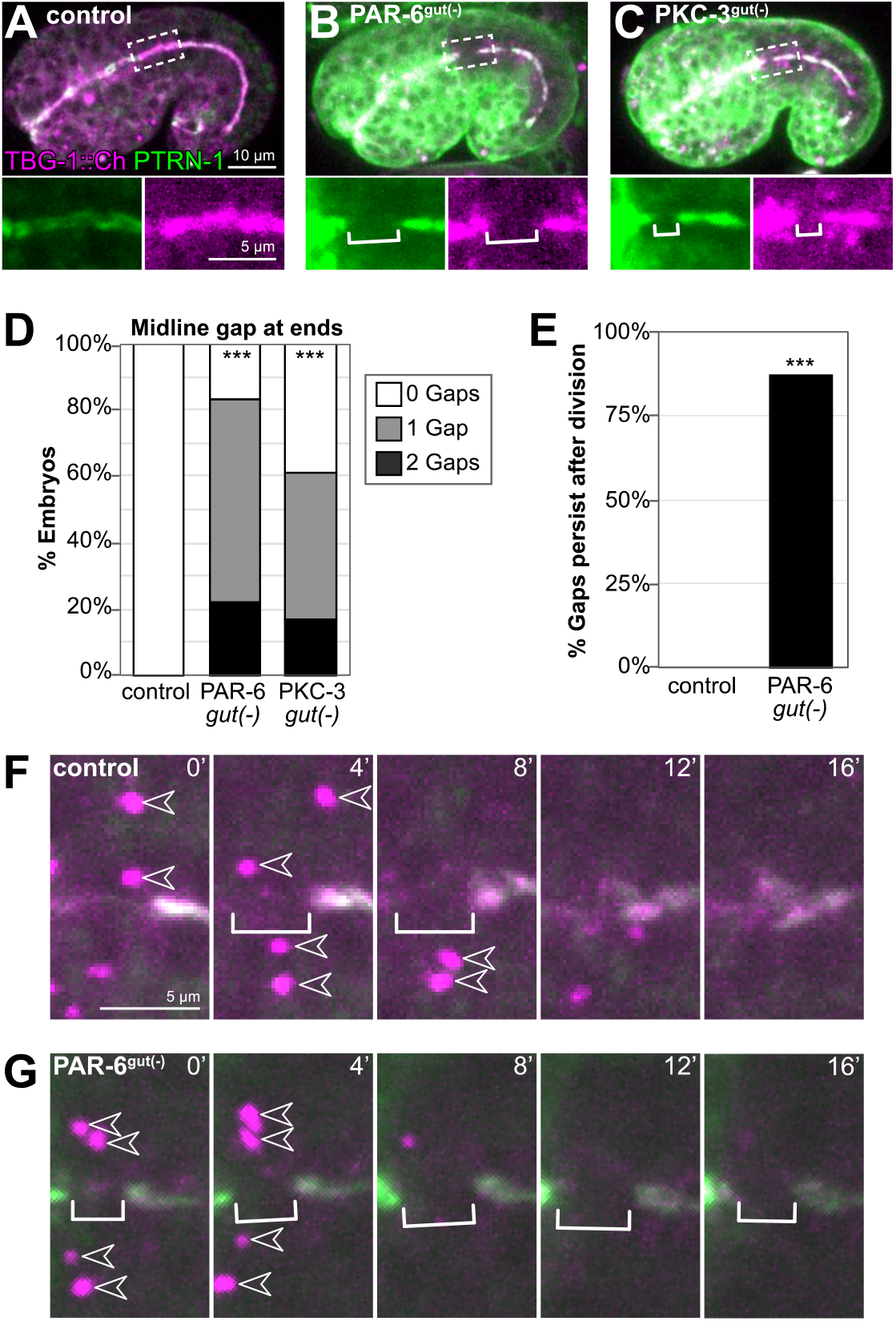
PAR-6 and PKC-3 are required for the return of MTOC function to the apical surface following mitosis. (A-C) Dorsolateral images of live comma- to 1.5-fold- stage embryos of indicated genotypes expressing transgenic MTOC markers TBG-1::mCherry (magenta) and endogenous PTRN-1::GFP (green). Maximum intensity Z-projections (1-3 microns) capture the intestinal midline. Enlarged images of boxed regions shown below. Star cell divisions had completed, but gaps (white bracket) were observed in PAR-6^gut(−)^ and PKC-3^gut(−)^ embryos. (B, C) Strong GFP fluorescence outside the intestine is undegraded PAR-6::ZF::GFP and ZF::GFP::PKC-3 in non-intestinal tissues. (D) Graph showing the percent of embryos with midline gaps in TBG-1 and PTRN-1 at the anterior and/or posterior regions of the intestine, approximately where star cell divisions occurred. Control: n = 22, PAR-6^gut(−)^: n = 18, PKC-3^gut(−)^: n = 18. Statistical analysis: ANOVA with Tukey’s post hoc tests. Differences from control indicated with asterisks. (E) Graph showing the percent of control (n=0/5) and PAR-6^gut(−)^ (n=7/8) embryos in which a midline gap in TBG-1 and PTRN-1 formed in synchronously dividing cells and persisted through the end of the 32-minute time lapse movie, with the gap itself persisting for at least 24 minutes. Statistical analysis: Fisher’s exact test. In control embryos, the gap lasted 8 minutes or less, except for one which lasted 12 minutes. (F, G) Live time-lapse imaging of TBG-1::mCherry and PTRN-1::GFP during star cell divisions in indicated genotypes. Bracket marks the apical MTOC gap; arrowheads indicate active centrosomal MTOCs marked with TBG-1 localization. All experiments used *ifb-2*p::*zif-1* to drive E8 onset of degradation. Scale bars = 10 μm in A, 5 μm in A inset and F. ***p<0.001.

In control embryos which had completed their star cell divisions (comma- to 1.8-fold stage), we observed continuous midline localization of the apical MTOC markers TBG-1::mCherry and PTRN-1::GFP (Fig. 2A). In contrast, we observed gaps in the apical MTOC in PAR-6^gut(−)^ and PKC-3^gut(−)^ embryos (Fig. 2B, C). Gaps were often observed toward the anterior and posterior ends of intestines, the approximate position of star cell daughters (Fig. 2D). To further test if gaps formed where star cells had divided, we used live time-lapse imaging of the MTOC in dividing star cells. No gaps were observed during asynchronous star cell divisions in control and PAR-6^gut(−)^ embryos (Supp. Fig. 3). In control embryos in which anterior star cells divided synchronously, a midline gap formed in the apical MTOC as centrosomes recruited additional TBG-1; MTOC proteins returned and filled the gap within 12 minutes as the centrosomes inactivated and shed TBG-1 (Fig. 2E, 2F). In PAR-6^gut(−)^ embryos, however, midline MTOC gaps that formed during synchronous divisions persisted long after mitotic exit through the end of the time course, for a minimum of 24 minutes (Fig. 2E, 2G). The persistence of the midline gap in MTOC proteins in PAR-6^gut(−)^ and PKC-3^gut(−)^ embryos is consistent with our hypothesis that PAR complex proteins promote the return of apical MTOC function after division.

**Figure 3.**
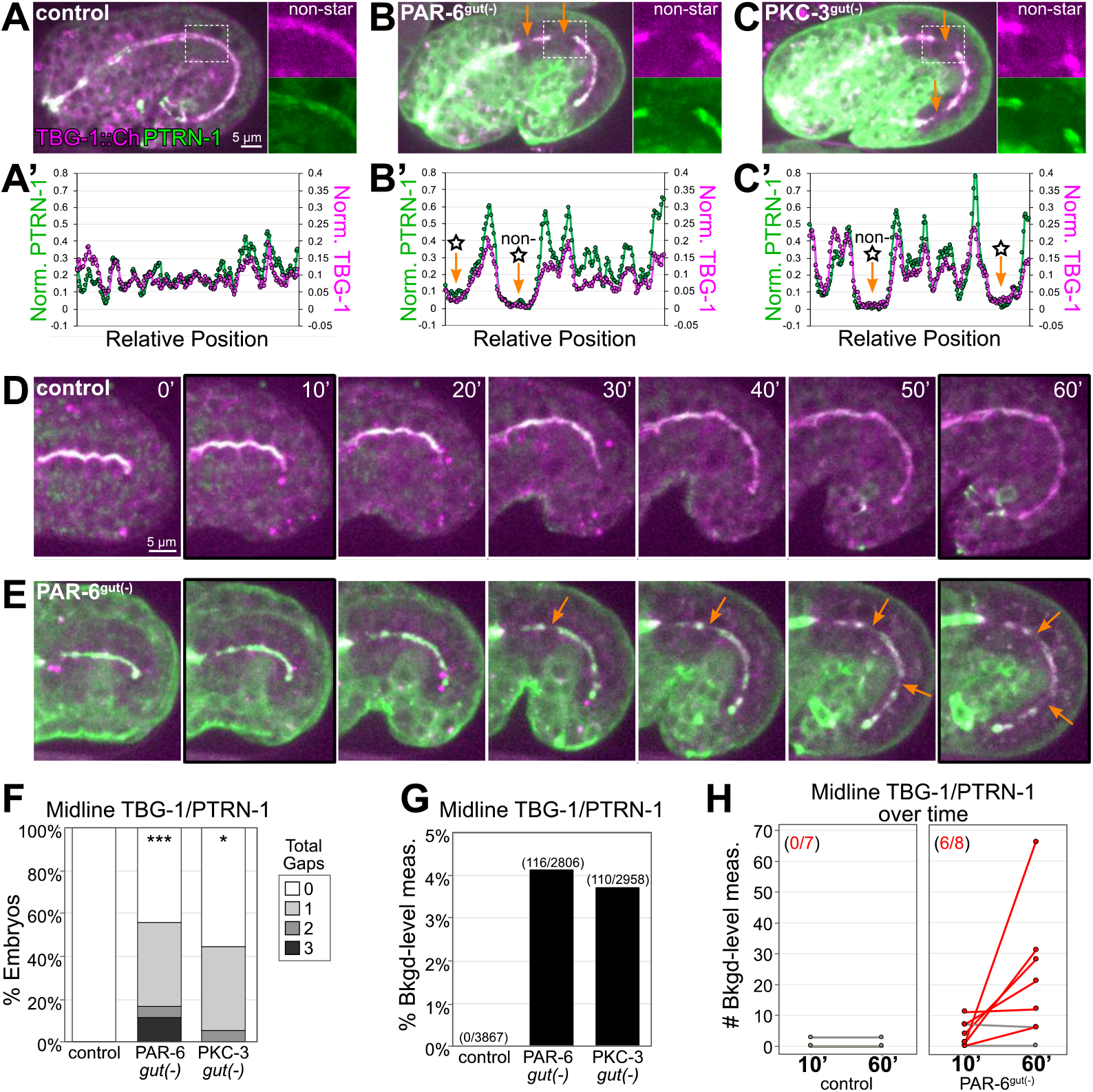
PAR-6 and PKC-3 are required to maintain a continuous apical MTOC during intestinal elongation. (A-C) Lateral live images of 1.5-1.8-fold-stage embryos of indicated genotypes expressing MTOC markers TBG-1::mCherry and PTRN-1::GFP. Maximum intensity Z-projections (2.5-3.5 microns) capture the intestinal midline. 2X magnified images of boxed region highlighting the apical MTOC in non-star cells are shown at right. Orange arrows indicate midline gaps. (A’-C’) Line scan along the apical midline of corresponding embryos above, plotting normalized PTRN- 1 or TBG-1 signal intensity from anterior to posterior. (D, E) Dorsal-dorsolateral view of posterior half of live embryos developing over 1 hour. Apical MTOC gaps are indicated (orange arrow). t=0’, the end of anterior star cell mitosis. (F) Graph of percent of embryos with midline gaps in MTOC proteins as defined by the normalized signal intensity of both TBG-1 and PTRN-1. See Methods for details. Control: n = 22, PAR-6^gut(−)^: n = 18, PKC-3^gut(−)^: n = 18. Statistical analysis: ANOVA with Tukey’s post hoc tests. Differences from control indicated with asterisks. No significant difference between PAR-6^gut(−)^ and PKC-3^gut(−)^. (G) Graph showing percent of midline intensity measurements in F that are “background-level.” Control: 0.00%, PAR-6^gut(−)^: 4.13%, PKC-3^gut(−)^: 3.72%. (H) Graph with paired comparisons per embryo in the number of background-level measurements at 10 minutes and 60 minutes after anterior star cell mitosis. Control: 0/7 embryos increased their number of background-level reads; PAR-6^gut(−)^: 6/8 embryos increased their number of background-level reads (red lines). All experiments used *ifb-2*p::*zif-1* to drive E8 onset of degradation. All experiments used *ifb-2*p::*zif-1* to drive E8 onset of degradation. Scale bars = 5 μm. *p<0.05, ***p<0.001.

### PAR-6 and PKC-3 are required during elongation to maintain apical MTOC continuity

In addition to the challenge that mitosis poses to the apical continuity of a polarized epithelium, the cell growth and shape changes underlying epithelial morphogenesis could similarly interrupt epithelial integrity. Intestinal length increases approximately 70% in the hour between the star cell divisions and 1.5-fold stage (from 22.4 ± 3.1 microns to 38.2 ± 2.2 microns), so we asked if PAR-6 and PKC-3 are required for the cell shape changes that the star cells and the non-dividing “non-star” cells undergo during intestinal elongation. We first assessed the continuity of the apical MTOC during elongation, using TBG-1::mCherry and PTRN-1::GFP as markers. In elongating control intestines, the apical MTOC remained continuous along the midline in both star cell daughters and in non-star cells (Fig. 3A, inset). A line scan tracing the midline of control intestines showed some variation in apical MTOC protein intensity but no apparent gaps in signal intensity (Fig. 3A’). By contrast, in elongating PAR-6^gut(−)^ and PKC-3^gut(−)^ intestines, midline gaps in MTOC proteins formed both in star cell daughters and in non-star cells that had not recently divided (Fig. 3B, 3C, insets); the midline gaps were clearly visualized by line scans (Fig. 3B’, 3C’). Consistent with elongation causing gaps to form, we observed only few PAR-6^gut(−)^ embryos with small midline gaps in TBG-1 and PTRN-1 prior to the star cell divisions and elongation (Control: n = 1/17, PAR-6^gut(−)^: n = 5/29). Using time-lapse imaging, we observed gaps in the apical MTOC emerge over time in individual embryos during elongation. In control intestines, the apical MTOC stayed continuous as the average midline length increased approximately 80% in 50 minutes (Fig. 3D). In PAR-6^gut(−)^ intestines, however, the apical MTOC formed gaps as the average apical surface elongated 70% in 50 minutes (Fig. 3E).

To quantify the apical MTOC defects in PAR-6^gut(−)^ and PKC-3^gut(−)^ embryos, we defined a “background-level” measurement as a midline intensity measurement for which both TBG-1::mCherry and PTRN-1::GFP signal intensities were in the same range as intestinal cytoplasmic background signal (see Methods), and a “gap” as three consecutive midline intensity measurements that were background-level. For this analysis, we included both star and non-star cell gaps. By these stringent criteria, 55.5% of PAR-6^gut(−)^ and 44.4% of PKC-3^gut(−)^ embryos had one or more gaps, compared with 0% of control embryos (Fig. 3F). As a total proportion of the midline MTOC line scan signal, we found that 4.1% of PAR-6^gut(−)^ and 3.7% of PKC-3^gut(−)^ midline measurements were background-level, compared to 0% in control intestines (Fig. 3G). The above criteria likely underestimate gap frequency, as more gaps were observed by eye (compare Fig. 3F to Fig. 4E). Finally, to determine if more gaps arose during elongation, we measured and compared MTOC protein signal intensity at the midline just after the anterior divisions complete (t=10’) and after 50 minutes of elongation (t=60’). The number of background-level measurements increased over time in PAR-6^gut(−)^ embryos but not in control embryos (Fig. 3H). The increased number of gaps in PAR-6^gut(−)^ embryos was likely not due to additional cell divisions, as we did not observe active centrosomes with TBG-1::mCherry, and intestines generally had 20 cells (n = 22/23); one embryo had 21 intestinal cells, as is occasionally observed in wild-type embryos (Sulston and Horvitz 1997), but it had no midline gaps. These results suggest that in addition to their roles in promoting continuity in dividing cells, PAR-6 and PKC-3 are also required for maintaining apical MTOC continuity between neighboring non-dividing cells during intestinal elongation.

**Figure 4.**
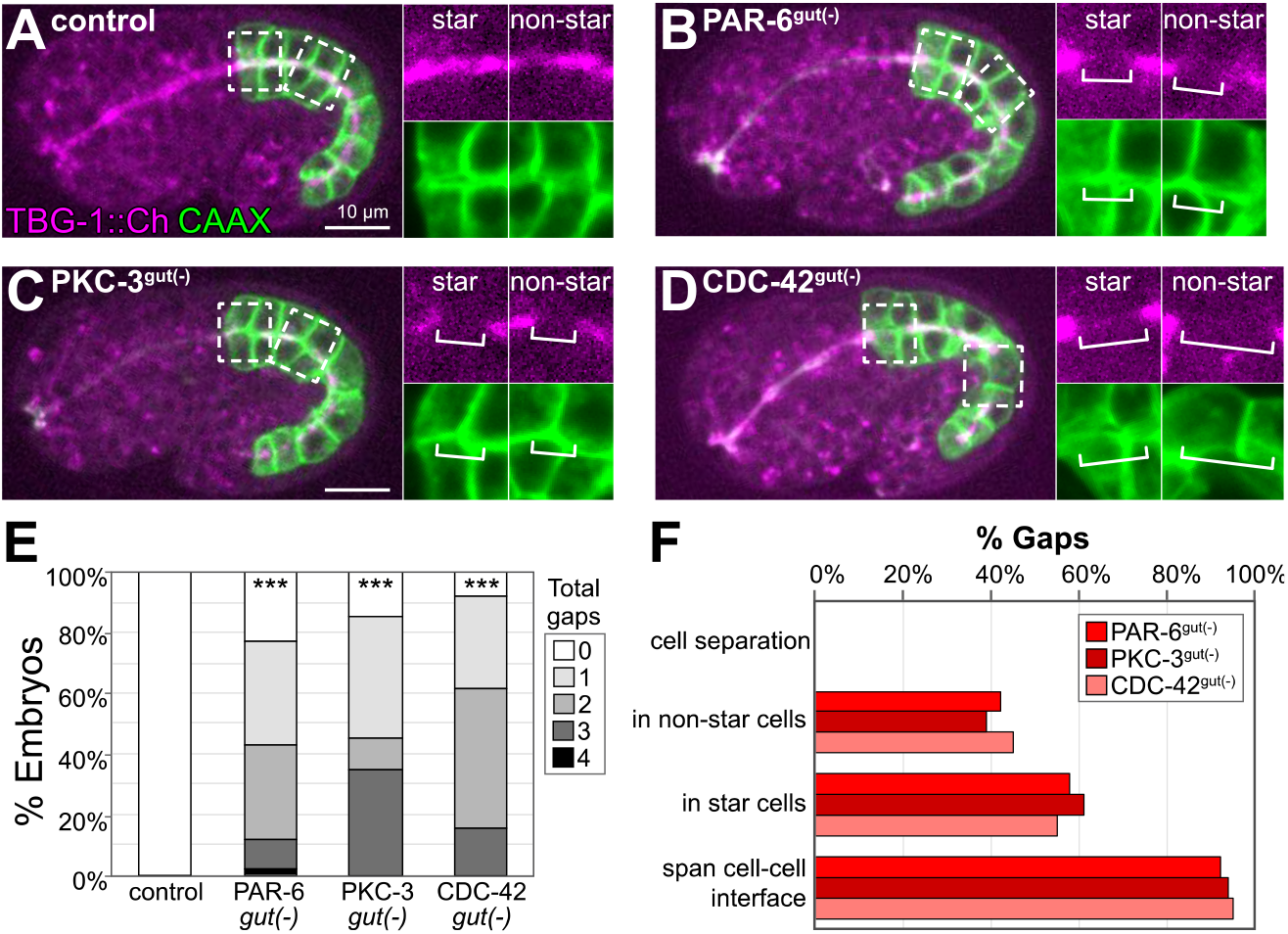
PAR-6, PKC-3, and CDC-42 are required for apical MTOC continuity but not for intestinal cell adhesion. (A-D) Lateral live images of 1.5-1.8-fold-stage embryos of indicated genotypes expressing intestine-specific GFP::CAAX and TBG-1::mCherry. Maximum intensity Z-projections (1-2 microns) capture the intestinal midline. 2X magnified images of boxed region highlighting the apical MTOC in star cells (left inset) and non-star cells (right inset). White brackets indicate midline gap. (E) Graph shows percent of embryos with indicated number of TBG-1 gaps; gaps were assessed by eye. Control: n = 41, PAR-6^gut(−)^: n = 44, PKC-3^gut(−)^: n = 20, and CDC-42^gut(−)^: n = 20. Statistical analysis: ANOVA with Tukey’s post hoc tests. Differences from control indicated with asterisks. No significant difference between the ^gut(−)^ strains. (F) Graph showing the percent of gaps of indicated genotypes that corresponded with physical detachment and separation of cells, that occurred in non-star cells versus star cell daughters, and that spanned the cell-cell interface between anterior/posterior neighbors. All experiments used *ifb-2*p::*zif-1* to drive E8 onset of degradation. Scale bars =10 μm. ***p<0.001.

### PAR-6, PKC-3, and CDC-42 depletion does not cause physical separation of intestinal cells

Previous studies demonstrated that PAR-6 is essential for correct junction formation in the intestine (Totong et al., 2007). Therefore, one possible cause of the midline gaps in MTOC proteins could be physical dissociation of intestinal cells due to defective cell adhesion. However, using TBG-1::mCherry to mark midline gaps and an intestine-specific GFP::CAAX to label cell membranes, we did not detect physical separation between intestinal cells in control, PAR-6^gut(−)^ or PKC-3^gut(−)^ embryos (Fig. 4A-C). The membrane marker clearly spanned the gaps in TBG-1 (Fig. 4A-C, insets), and neighboring intestinal cells remained adjacent to each other.

PKC-3 kinase activity can be activated by the Rho GTPase CDC-42 in many epithelial contexts (Rodriguez-Boulan and Macara, 2014). If PKC-3 kinase activity is important for maintaining apical continuity, we predicted that CDC-42 would also be important and that depleting CDC-42 in the intestine would cause similar gap defects. Indeed, like PAR-6 ^gut(−)^ and PKC-3 ^gut(−)^ intestines, CDC-42^gut(−)^ intestines had midline gaps in TBG-1 in similar numbers and positions (Fig. 4D-F).

### Midline gaps in the apical MTOC also lack apical and junctional proteins and fail to exclude basolateral proteins

To determine if the midline gaps in the MTOC were specific to MTOC proteins, or if they reflected more general discontinuities in the apical domain, we examined the localization of other apical markers. In PAR-6^gut(−)^ embryos, apical YFP::ACT-5/actin formed gaps along the midline in both star cell daughters and in non-star cells (10/19 and 9/19 gaps, respectively; Fig. 5A, B), similar to MTOC protein gaps (Fig. 4F). In addition, we assessed localization of PAR-3, the most upstream apical marker known in the intestine (Achilleos et al., 2010), and found midline gaps in PAR-3::tagRFP that overlapped with gaps in PTRN-1::GFP (Fig. 5C, F). These results suggest that PAR-6 is essential not only for a continuous apical MTOC but also for the general continuity of the apical domain during elongation.

**Figure 5.**
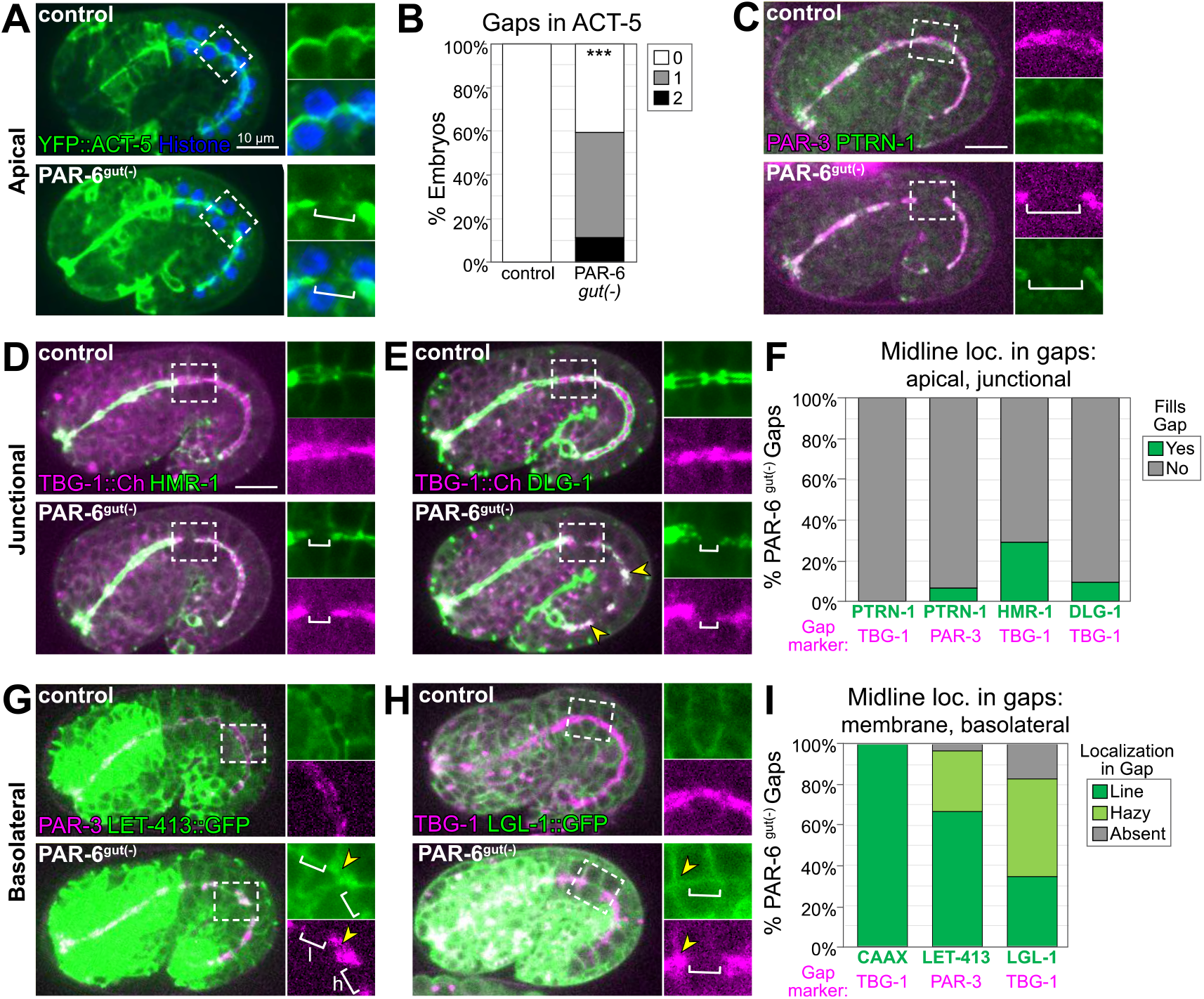
Midline gaps in apical and junction proteins overlap and fail to exclude basolateral proteins in PAR-6^gut(−)^ embryos. (A-H) Lateral live images of 1.5-1.8-fold-stage embryos of indicated genotypes expressing indicated markers. Maximum intensity Z-projections capture the intestinal midline, except LGL-1 which is a single Z-slice. 2X magnified images of boxed region highlighting midline gaps are shown at right. White brackets indicate midline gap. Yellow arrowheads indicate colocalization of indicated markers. Note that DLG-1::GFP contrast and brightness in (E) was increased in inset PAR-6^gut(−)^ image to visualize gap. (B) Graph shows percent of embryos with indicated number of ACT-5 gaps. Control n = 19 embryos, PAR-6^gut(−)^ n = 27. Statistical analysis: Student’s t-test. (F) Graph showing the percent of midline gaps (magenta) in PAR-6^gut(−)^ embryos to which the indicated protein (green) localized. PTRN-1/TBG-1: n = 23 gaps, 18 embryos; PTRN-1/PAR-3: n = 86 gaps, 35 embryos; HMR-1/TBG-1: n = 31 gaps, 37 embryos; DLG-1/TBG-1: n = 31 gaps, 23 embryos. (G, H) LET-413::GFP and LGL-1::GFP localization could appear as a line (G inset, “l”) or as a hazy enrichment (“h”). (I) Graph showing the percent of midline gaps (magenta) to which the indicated protein (green) localized in PAR-6^gut(−)^ intestines. Midline gap localization of LET-413 and LGL-1 was scored only for anterior cells (int1-4). CAAX/TBG-1: n = 49 gaps, 36 embryos; LET-413/PAR-3: n = 30 anterior gaps, 18 embryos; LGL-1/TBG-1: n = 29 anterior gaps, 45 embryos. All experiments used *ifb-2*p::*zif-1* to drive E8 onset of degradation except A-C, which used *elt-2*p::*zif-1* to drive degradation at E4. Scale bars = 10 μm. ***p<0.001.

Our analysis of GFP::CAAX in PAR-6^gut(−)^ embryos revealed that 92% of apical gaps spanned the interface between adjacent cells (Fig. 4F). We hypothesized that junctional or basolateral proteins might be inappropriately invading and spreading into the apical surface, creating the observed gaps in the apical domain. To test this hypothesis, we examined the localization of two apical junction complex proteins, HMR-1/E-cadherin and DLG-1/Dlg (Bossinger et al., 2001; Costa et al., 1998), relative to gaps in TBG-1::mCherry in PAR-6^gut(−)^ embryos. Normally, HMR-1 and DLG-1 localize as a continuous band at the cell periphery between the apical and basolateral domains and form ladder-like belt junctions at the interfaces between intestinal cells ((Leung et al., 1999; McMahon et al., 2001), Supp. Fig. 1). In PAR-6^gut(−)^ embryos, HMR-1::GFP localized to the outer edge of the TBG-1 signal as expected for a junctional protein, but was generally absent from the TBG-1 midline gaps (Fig. 5D, F). For small TBG-1 gaps, a gap in HMR-1 was sometimes not observed. We also examined DLG-1::GFP localization, which was severely perturbed along the entire midline in PAR-6^gut(−)^ embryos as has previously been reported following PAR-6 depletion (Totong et al., 2007). We occasionally found large patches of DLG-1 that colocalized with TBG-1; however, DLG-1 was clearly absent from the majority of apical gaps (Fig. 5E, F). Together, these results indicate that junctional complexes do not spread into and fill gaps in apical proteins following PAR-6 depletion.

Lastly, we asked whether the basolateral domain expanded and spread into midline gaps in PAR-6^gut(−)^ embryos. We examined two basolateral markers: LET-413/Scribble, which localizes to basolateral and junctional domains (Legouis et al., 2000; McMahon et al., 2001), and LGL-1, which localizes specifically to basolateral domains (Beatty et al., 2010). Because of a natural twist in the intestine (Hermann et al., 2000; Leung et al., 1999), the apical domain in anterior cells presented itself as a side view under normal imaging conditions, making it easier to detect the localization of membrane-enriched proteins at the midline of these cells (compare midline GFP::CAAX in anterior and posterior cells of middle image in Fig. 1C). Therefore, we restricted our analysis of LET-413 and LGL-1 localization at TBG-1::mCherry gaps to the anterior half of the intestine (int1-4). We did not observe LET-413::GFP (n=18/20) or LGL-1::GFP (n=18/19) localization at the apical surface in control intestines. In most anterior gaps in TBG-1 in PAR-6^gut(−)^ embryos, LET-413 localization appeared as a line (Fig. 5G inset “l”, 5I), with some appearing as a hazy enrichment (Fig. 5G inset “h”). The lines of LET-413 in the apical gaps likely reflect the mislocalization of basolateral rather than junctional LET-413 as the other junctional proteins we examined were absent at apical gaps. Consistent with this interpretation, we also frequently saw LGL-1 localize to the midline gaps either as a line or as a hazy enrichment (Fig. 5H, I). However, we also observed stretches of the midline in PAR-6^gut(−)^ embryos where LET-413 or LGL-1 colocalized with the apical marker (Fig. 5G, H, inset yellow arrowhead), suggesting a failure of LET-413 and LGL-1 exclusion from the apical surface rather than specific invasion of the basolateral domain into the midline gaps.

### PAR-6^gut(−)^ and PKC-3^gut(−^intestinal lumens are functionally obstructed and cannot pass food

The importance of PAR-6 for building a functional intestine is unknown, because global loss of *par-6* causes lethality in embryos, well before intestinal function can be assessed (Totong et al., 2007; Watts et al., 1996). Our tissue-specific depletion strategy thus enabled us to bypass *par-6* embryonic lethality and examine intestinal morphology and function in PAR-6^gut(−)^ larvae. We hypothesized that defects in apical and junctional continuity in embryonic intestines would lead to morphological and functional defects in the larval intestine. Indeed, all PAR-6^gut(−)^ and PKC-3^gut(−)^ larvae arrested as young L1 larvae (Fig. 6A, Supp. Table 1), suggesting that intestinal function was severely compromised. Surprisingly, only 39% of CDC-42^gut(−)^ larvae arrested as L1 larvae.

**Figure 6.**
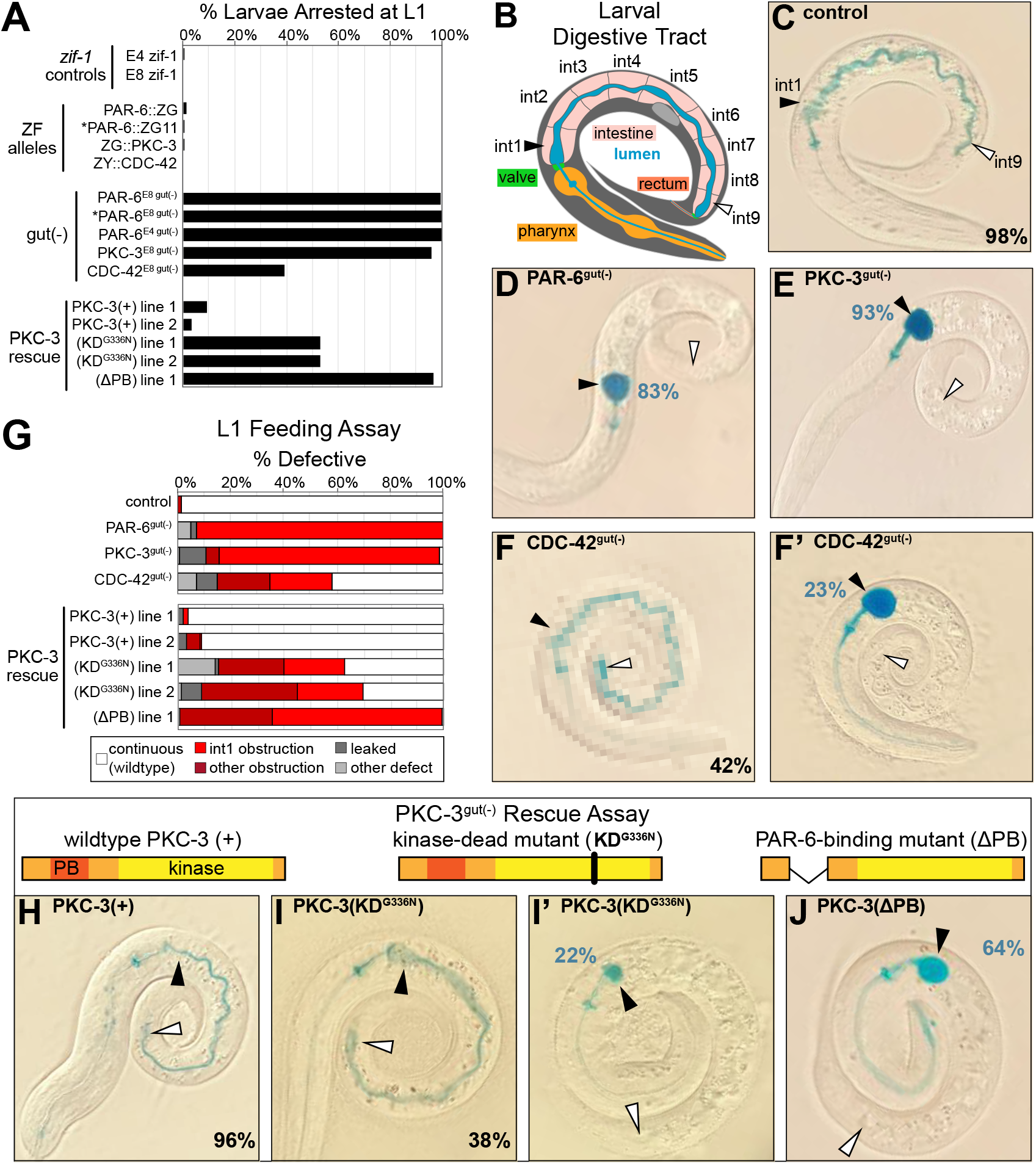
PAR-6, PKC-3, and CDC-42 are essential in the embryonic intestine for larval viability and intestinal function. (A) Graph of the average percent of larvae from three trials that arrested in the L1 stage 72 hours after embryos were laid (see Supp. Table 1). Genotypes as indicated. Asterisks mark the PAR-6::ZF::GFP_11_ allele (See Methods). For PKC-3 rescue, PKC-3^gut(−)^ larvae carried a transgene providing intestine-specific expression of wild-type or mutant PKC-3. Each line was independently isolated. (B) Cartoon schematic of the larval digestive tract and the path of blue-dyed food in the feeding assay in C-J. (C-J) Color images of L1 larvae of indicated genotypes after 3 hour incubation in blue-dyed food. Arrowheads indicate the positions of first intestinal ring int1 (black) and the last ring int9 (white). Percentages indicate frequency of continuous lumen (C, F, H, I) and int1 obstruction (D, E, F’, I’, J). (G) Graph showing the average percent of L1 larvae from three trials that had defective lumens (see Supp. Table 2). (H-J) Cartoon schematic of PKC-3 mutations shown above the corresponding PKC-3^gut(−)^ larvae carrying a wild-type or mutant PKC-3 transgene. All experiments used *ifb-2*p::*zif-1* to drive E8 onset of degradation.

To better understand the larval arrest caused by intestine-specific loss of these polarity regulators, we asked if intestinal barrier function was disrupted, or if the lumen was discontinuous and thus obstructed. We used a “Smurf” feeding assay to distinguish between these two possibilities ((Gelino et al., 2016; Rera et al., 2011), Fig. 6B; Supp. Table 2), feeding worms blue-dyed food to monitor whether it leaked into the body (barrier dysfunction) or was trapped in the intestinal lumen (obstructed / discontinuous lumen). In 98% of control larvae, blue food traveled through and filled the entire lumen without leaking into the body (Fig. 6C, G). However, in 83% of PAR-6^gut(−)^ larvae and 93% of PKC-3^gut(−)^ larvae, blue food passed through the pharynx and pharyngeal valve but was trapped at the anterior end of the intestine, suggesting the lumen in the first intestinal ring (int1) was obstructed (Fig. 6D, E, G). Blue food leaked into the body of a small percentage of larvae (10% of PAR-6^gut(−)^, 2% of PKC-3^gut(−)^), suggesting that most of the PAR-6^gut(−)^ and PKC-3^gut(−)^ larvae that ingested food retained their intestinal barrier function, at least between the pharyngeal valve and int1. Consistent with their milder larval arrest phenotype, CDC-42^gut(−)^ larvae also showed a lower penetrance of obstructed intestines; 42% had a normal intestine (Fig. 6F), 23% had an early int1 obstruction (Fig. 6F’), and 20% had a more posterior obstruction.

The interdependent localization of PAR-6 and PKC-3 obscures which of their respective scaffolding and kinase functions is important for maintaining apical MTOC continuity and building a functional intestine. To determine if PKC-3 kinase activity is required to build a functional intestine, we performed a rescue assay with three forms of PKC-3: wild-type PKC-3(+); PKC-3(∆PB), which removes the PAR-6-binding domain (Kim et al., 2009); and PKC-3(G336N), for which the analogous mutation in *Drosophila* aPkc abolishes *in vitro* kinase activity to the same degree as the canonical kinase-dead mutant K293A without losing Par6 binding activity (Kim et al., 2009). We found that wild-type PKC-3 was able to rescue both the larval arrest and obstructed lumen defects (Fig. 6A, G, H); the kinase mutant showed variable rescuing activity (Fig. 6A, G, I, I’); and the PAR-6-binding mutant showed almost no rescuing activity (Fig. 6A, G, J). The PKC-3 kinase mutant rescue showed striking similarity to the CDC-42^gut(−)^ phenotype.

### PAR-6^gut(−)^ and PKC-3^gut(−)^ intestinal lumens have edematous swellings and multiple constrictions

The intestinal obstructions in PAR-6^gut(−)^ and PKC-3^gut(−)^ larvae could be caused by constricted or closed lumens, so we next examined cell arrangement and tissue morphology with a membrane marker (intestine-specific GFP::CAAX) and DIC imaging. Unlike the continuous open lumen of control intestines (Fig. 7A), the lumens of PAR-6^gut(−)^ and PKC-3^gut(−)^ intestines frequently appeared to be pinched shut in multiple places, often with edematous swellings (Fig. 7B, C, E). CDC-42^gut(−)^ larvae generally had one or two points of luminal constriction (Fig. 7D), but the overall intestinal morphology was not as severely defective as PAR-6^gut(−)^ and PKC-3^gut(−)^ larvae. Based on the intercellular position of the midline gaps in embryonic intestines depleted of polarity regulators, we predicted that lumen formation would frequently fail at the interface between adjacent int rings. While control intestines clearly had an open lumen between int rings, PAR-6^gut(−)^ and PKC-3^gut(−)^ intestines all had one or more int interface with a constricted lumen, and CDC-42^gut(−)^ larvae again showed a milder phenotype (Fig. 7F). Further, most PAR-6^gut(−)^ and PKC-3^gut(−)^ larvae and 50% of CDC-42^gut(−)^ larvae had their lumens pinched shut within int1 (Fig. 7G), consistent with the int1 obstruction and accumulation of blue-dyed food in the feeding assay and with the midline gaps that formed in embryos after the anterior star cell divisions. Thus, PAR-6 and PKC-3 are essential during embryonic development to build an intestine with a continuous open lumen.

**Figure 7.**
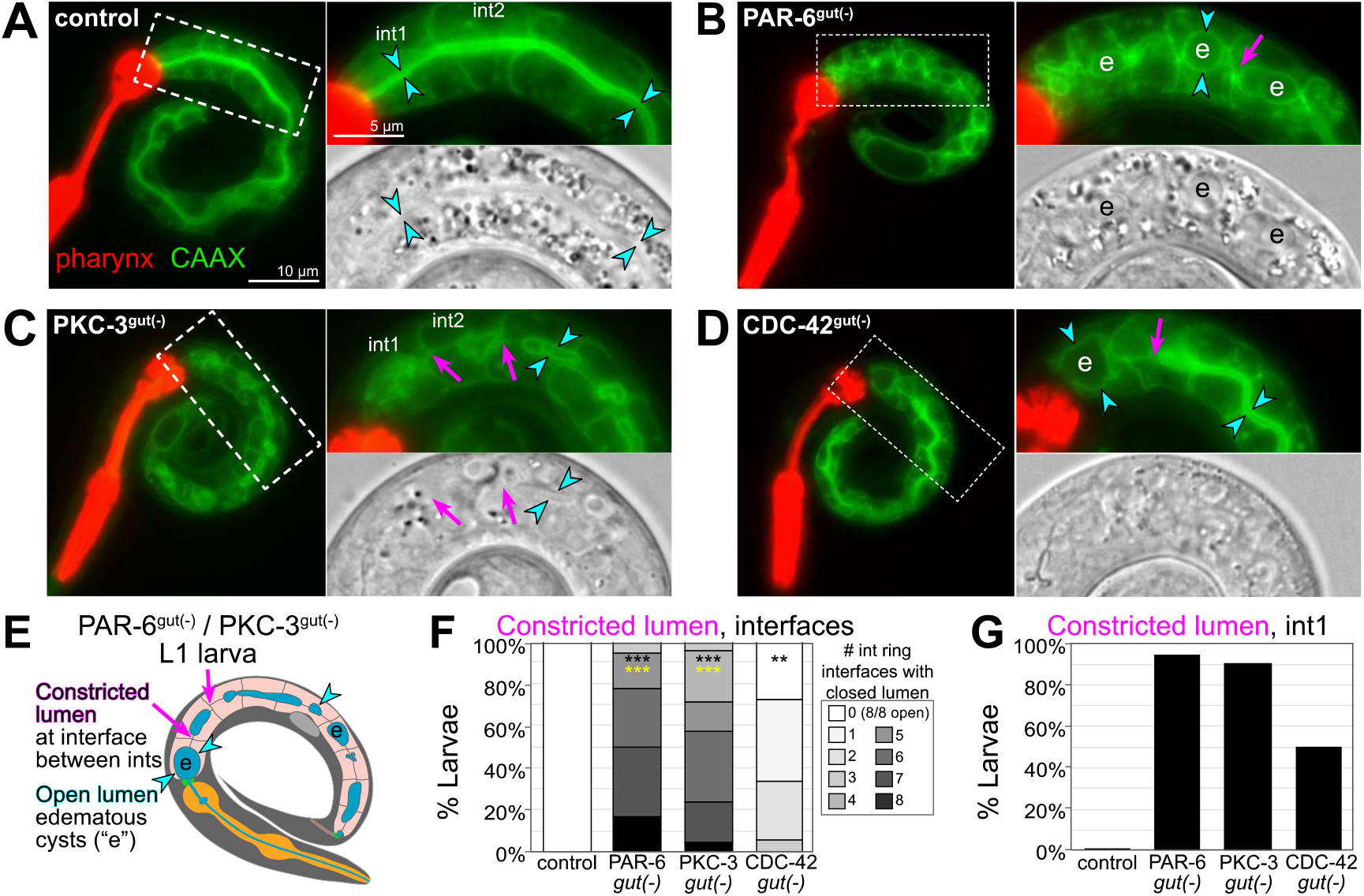
PAR-6, PKC-3, and CDC-42 are essential in the embryonic intestine for larval intestine morphology. (A-D) Live images of L1 larvae of indicated genotypes with GFP::CAAX-labeled intestinal membranes and mCherry-labeled pharynxes. Maximum intensity Z-projections (0-1 microns) capture the intestinal midline. 2X magnified images of boxed region on the right; GFP::CAAX (top) and DIC micrograph (bottom). Intestinal lumen (paired cyan arrowheads), edematous luminal swellings (“e”), and constricted lumens (magenta arrows) are indicated. (E) Cartoon model of the defects in PAR-6^gut(−)^ and PKC-3^gut(−)^ intestines. (F) Graph showing percent of larvae with the indicated number of int ring interfaces with a constricted lumen (see magenta arrows in E). Control: n = 20, PAR-6^gut(−)^: n = 18, PKC-3^gut(−)^: n = 18, and CDC-42^gut(−)^: n = 21. Statistical analysis: ANOVA with Tukey’s post hoc tests. Differences from control indicated with black asterisks. Differences from CDC-42^gut(−)^ indicated with yellow asterisks. No significant difference between PAR-6^gut(−)^ and PKC-3^gut(−)^. (G) Graph showing percent of larvae with a constricted lumen in int1. All experiments used *ifb-2*p::*zif-1* to drive E8 onset of degradation. Scale bars = 10 μm in A-D, and 5 μm in insets. **p<0.01, ***p<0.001.

### Apical and junctional proteins are discontinuous in larval PAR-6^gut(−)^ intestines

In examining the membranes of PAR-6^gut(−)^ intestinal cells, the edematous swellings nearly always occurred along the midline, where an apical surface and lumen would normally be built. To further understand how the early gap defects correlated with these later morphological defects, we examined the apical markers PAR-3::GFP and YFP::ACT-5, the junctional marker DLG-1::GFP, and the basolateral marker LGL-1::GFP, which remained expressed in L1 larvae. Unlike the continuous luminal PAR-3 in control L1 larval intestines, PAR-3 localized to hollow sphere-like structures that were often separated by gaps in localization in PAR-6^gut(−)^ larvae (Fig. 8A-C), consistent with a discontinuous luminal surface. ACT-5 is strongly expressed in larval intestines, and unlike the continuous luminal ACT-5 in control larvae, all PAR-6^gut(−)^ larvae showed multiple gaps in luminal ACT-5 (Fig. 7D-F). We also found DLG-1 localization to be discontinuous. While control intestines contained a continuous ladder-like belt arrangement of DLG-1, all PAR-6^gut(−)^ larvae had DLG-1 bands that were clearly discontinuous between adjacent int rings in multiple places (Fig. 8G-I). Lastly, we examined LGL-1 localization to determine if basolateral proteins continued to mislocalize to the larval intestinal midline. In control larvae, LGL-1 was clearly excluded from the apical surface, but in many PAR-6^gut(−)^ larvae, we observed localization of LGL-1 to the midline (Fig. 8J-L), suggesting a continued failure to exclude basolateral proteins in PAR-6^gut(−)^ larvae. Together, these results suggest that the early gaps in apical and junctional proteins and the failure to exclude basolateral proteins at the midline were retained during development, leading to intestines with discontinuous apical surfaces and junctions separated by midline regions where lumen formation did not occur.

**Figure 8.**
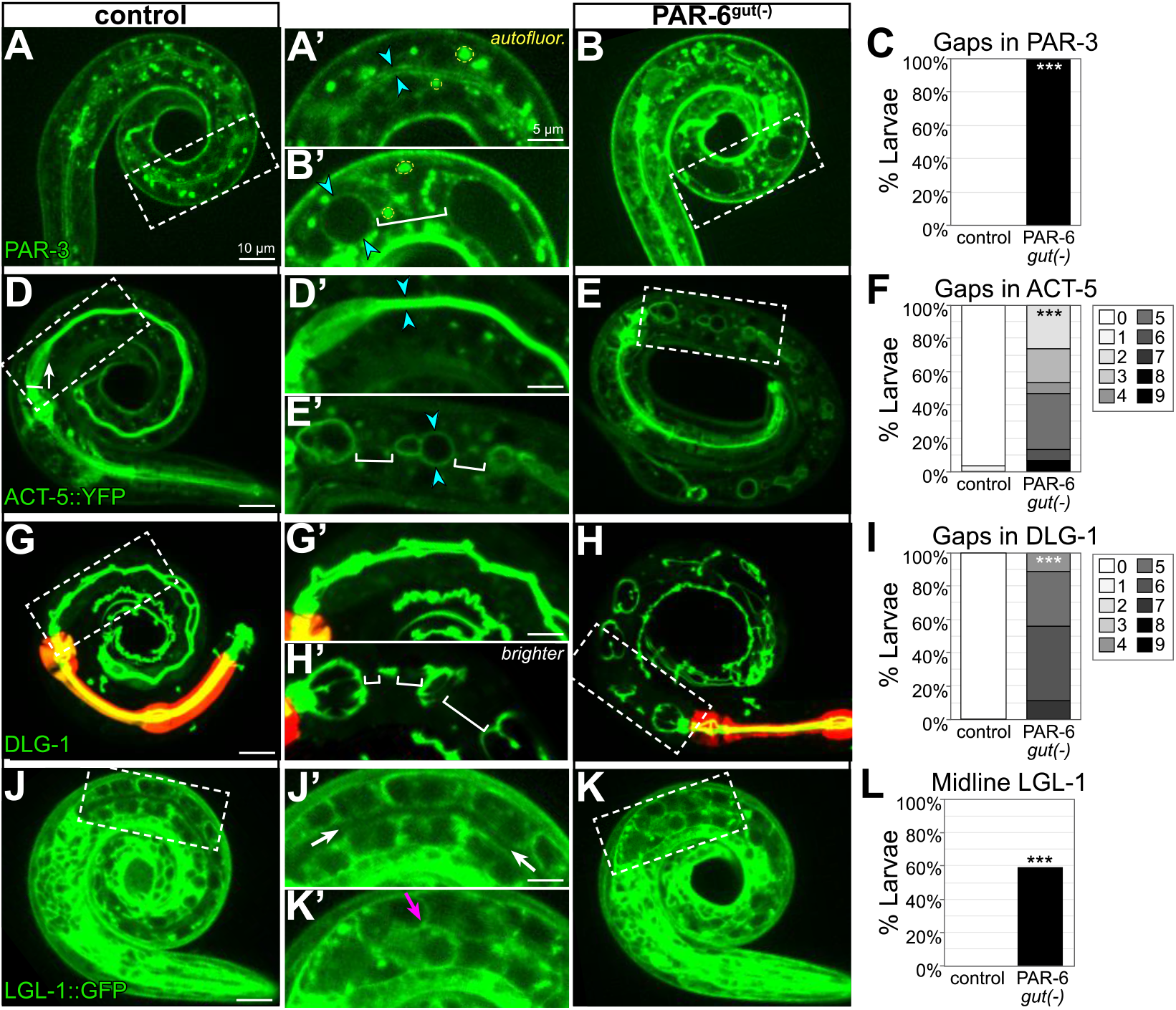
PAR-6^gut(−)^ larval intestines have gaps in apical and junctional proteins and a discontinuous lumen. Live images of L1 control and PAR-6^gut(−)^ larvae expressing the indicated marker, with adjacent 2X magnified images of boxed region, and graphs quantifying luminal localization defects. Maximum intensity Z-projections capture the intestinal midline for all images except for the minimum-intensity projections of LGL-1::GFP that better visualize the absence of apical LGL-1 in control larvae. Open intestinal lumens (cyan arrowheads), gaps in protein localization (brackets), midline protein localization presence (magenta arrows) or absence (white arrows) are indicated. Yellow dashed circles outline examples of the bright autofluorescent puncta from birefringent gut granules. (C) Percent of L1 larvae with any gaps in apical PAR-3::GFP. Control: n = 16, PAR-6^gut(−)^: n = 21. (F) Percent of L1 larvae with indicated number of gaps in apical YFP::ACT-5. Control: n = 29, PAR-6^gut(−)^: n = 15. (I) Percent of larvae with indicated number of gaps in DLG-1. Control: n = 23, PAR-6^gut(−)^: n = 18. (L) Percent of L1 larvae with any midline-localized LGL-1. Control: n = 15, PAR-6^gut(−)^: n = 27. All experiments used *ifb-2*p::*zif-1* to drive E8 onset of degradation except D-F, which used *elt-2*p::*zif-1* to drive degradation at E4. Scale bars = 10 μm. Statistical analyses: Student’s t-test (F,I) and Fisher’s exact test (C, L). ***p<0.001.

## DISCUSSION

PAR complex proteins are essential, highly conserved polarity regulators that play critical roles in epithelial cell polarity establishment and for apical and junctional maturation and maintenance. It is often difficult to tease apart their requirement for these different roles, especially *in vivo*, and to determine how morphological defects caused by their removal during development affect the mature organ. Using tissue-specific protein degradation to deplete proteins from developing intestines, we found that the PAR complex proteins PAR-6, PKC-3, and CDC-42 are required to preserve apical and junctional continuity as cells in the developing intestine divide and elongate during morphogenesis. This continuity is critical for transforming the polarized intestinal primordium into a functional tube at hatching, as depletion of any of these proteins resulted in a failure to build a continuous lumen that could pass food. Taken together, our data suggest a model in which PAR-6, PKC-3, and CDC-42 promote apical, junctional, and basolateral domain remodeling to maintain tissue-level continuity during intestinal cell division and elongation, thereby ensuring the formation of an open and functional intestinal lumen (Fig. 9).

**Figure 9.**
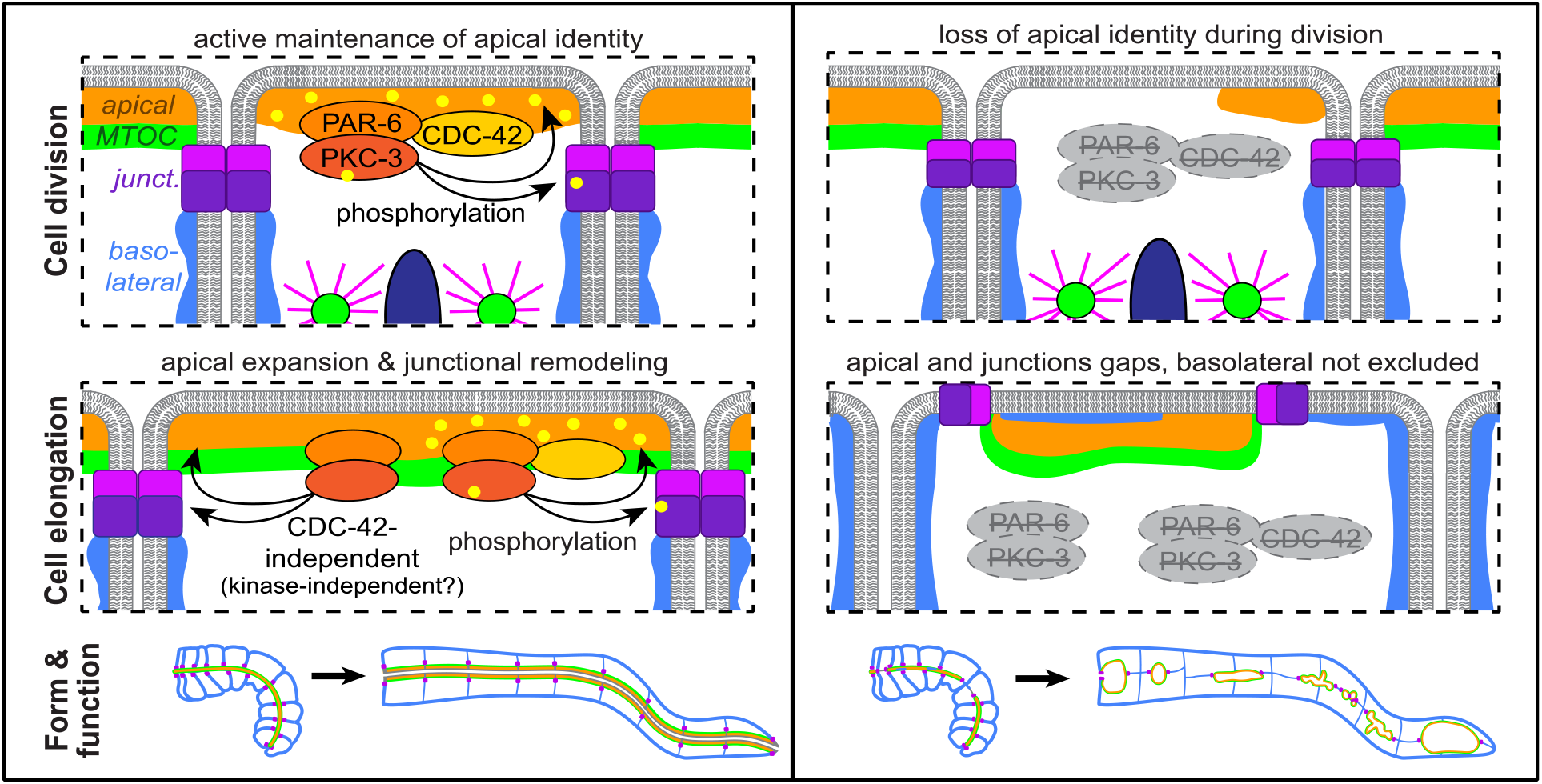
Model. A cartoon model of the role of PAR-6, PKC-3, and CDC-42 in apical and junctional remodeling as *C. elegans*

### The contribution of tissue-specific depletion studies

A critical feature of this study was our tissue-specific protein degradation approach. Ubiquitous somatic depletion of PAR-6, PKC-3, or CDC-42 causes embryonic lethality and failed elongation due to skin defects (Montoyo-Rosario et al., 2020; Totong et al., 2007; Zilberman et al., 2017), thereby masking potential roles of these proteins in other tissues during or after elongation. Using tissue-specific degradation, we bypassed the requirement for these proteins in earlier embryonic development and in other tissues, and revealed the requirement for PAR-6, PKC-3, and CDC-42 for apical and junctional continuity during intestinal morphogenesis. While we found that PAR-6 and PKC-3 were required embryonically to build a functional intestine, recently PAR-6 and PKC-3 were found to be dispensable in the larval intestine, even though intestinal elongation continues through larval development (Castiglioni et al.). These findings underscore another advantage of tissue-specific protein depletion: we can control the timing of depletion to uncouple early and late roles in organ development and maintenance. Thus, we can determine that PAR-6 and PKC-3 are required during embryonic morphogenesis to build the intestine, but that larvae hatch and use a different mechanism to continue to elongate the intestine.

### PAR complex protein requirement during epithelial cell mitosis

Our study found that dividing cells were sensitive to depletion of PAR-6, PKC-3 and CDC-42 during intestinal morphogenesis. Star cells contribute only three of the nine int rings that build the intestine (Supp. Fig. 1), yet over half of the midline gaps in 1.5-fold embryos affected star cell daughters. Dividing epithelial cells extensively remodel many polarized features, any of which could depend on PAR complex proteins. The apical MTOC and microtubules are removed and centrosomes are reactivated to organize microtubules into a bipolar spindle. Apical microtubules contribute to the establishment and maintenance of apical and junctional domains (Feldman and Priess, 2012; Harris and Peifer, 2005; Vasileva and Citi, 2018), so losing apical microtubules during mitosis could weaken the integrity of an epithelial cell and make it more dependent on the PAR complex to maintain polarity. In *Drosophila* neuroblasts, parallel inputs from PAR complex proteins and microtubules assemble a polarized Pins/Gαi crescent to orient the mitotic spindle (Siegrist and Doe, 2005). Similarly, apical microtubules and PAR-6 could redundantly maintain the apical domain in star cells; loss of the apical domain would thus only occur when both apical microtubules and PAR-6 are absent, as in dividing PAR-6^gut(−)^ star cells. As microtubule orientation is opposite in these two cases, each context would likely use different motor proteins. For example, intestinal star cells could use dynein to transport vesicles that replenish and maintain the apical domain and junctions (Rodriguez-Boulan and Macara, 2014).

Division also requires the formation of new junctions between daughter cells (Baum and Georgiou, 2011; Roignot et al., 2013), a process that could specifically rely on the known ability of PAR complex proteins to regulate junction formation and maturation during polarity establishment. The adherens junctions of interphase cells can promote tissue integrity in dividing neighbor cells by orienting the mitotic spindle, maintaining adhesion, and regulating the length of the new cell-cell interface (Gloerich et al., 2017; Herszterg et al., 2013; Higashi et al., 2016). Synchronous division of neighboring cells therefore creates a new challenge as both cells remodel their junctions at the same time, which could temporarily remove important positional information and adhesion. Indeed, we observed that when wild-type star cells divided synchronously, HMR-1 and DLG-1 levels were drastically reduced or absent. In addition, when PAR-6^gut(−)^ star cells divided synchronously, persistent midline gaps in the apical MTOC formed, but we did not observe such gaps during asynchronous divisions. These results suggest that either apical PAR-6 or a non-dividing neighboring cell is sufficient for new daughter cells to correctly remodel and build junctions, but that loss of both can impair this remodeling. Thus, adherens junctions may function as an external orientation cue to ensure correct polarity and junction position in daughter cells.

### Gap formation during epithelial cell elongation – cause versus consequence

Elongating epithelial tubes often must increase the area of the apical domains and junctions of their underlying cells, and consistently, the midline gap phenotype in PAR-6^gut(−)^ and PKC-3^gut(−)^ embryos became more severe during the greater than three-fold increase in intestine length that occurred by the L1 larval stage. Most gaps occurred between anterior and posterior cells without physical cell separation, raising the question of what causes cell-cell interfaces to be vulnerable to loss of PAR-6 or PKC-3. One possibility is that weakened junctions allow basolateral domains to invade and compress the apical domain. While we observed some basolateral protein localization in the gaps in PAR-6^gut(−)^ embryos, the signal was relatively weak and was not specific to the gaps, indicating that basolateral protein localization to the midline does not create apical and junctional gaps. Alternatively, compromised apical and junctional expansion could allow basolateral proteins to diffuse into the new midline space created by elongation. Apical expansion could fail if, for example, the mobility of important apical determinants like PAR-3 decreased and failed to spread to newly added midline membrane. Membrane lacking PAR-3 would likely have less PI(4,5)P_2_ and no apical microtubules, both of which normally promote apical protein localization and recruitment (Campanale et al., 2017; Rodriguez-Boulan and Macara, 2014). Failure in junctional remodeling could also impede apical expansion by creating junctions that corral and trap the apical domain as cells elongate. Positioning junctions, apical, and basolateral domains is by necessity a highly intertwined process, as each domain helps reinforce the position of the other, so it is difficult to draw firm conclusions about the causes of the gaps versus the consequences.

### Embryonic PAR complex proteins and intestinal tissue integrity

The primary defect in PAR-6-, PKC-3-, and CDC-42-depleted intestines is a failure to maintain the coordination of apicobasal polarity between cells; apical and junctional gaps correlated with luminal gaps in larval intestines, the luminal gaps blocked the passage of food, and young larvae arrested and died. Despite the lethality, many aspects of epithelial tissue integrity appeared intact in these larval intestines and thus did not require PAR-6, PKC-3, or CDC-42.

#### Selective barrier function

Disrupting tight/septate junction proteins can lead to defects in epithelial barrier function and thus loss of tissue integrity (Asano et al., 2003; Behr et al., 2003). Intestines depleted of PAR-6, PKC-3 or CDC-42 rarely leaked, so barrier function appeared largely intact. This result was unexpected due to the known requirement for PAR-6 in junction maturation (Totong et al., 2007). While junctional DLG-1 localization was dim and fragmented in PAR-6^gut(−)^ embryos ((Totong et al., 2007), this study), DLG-1 appeared to recover a junction-like localization in PAR-6^gut(−)^ larvae. In addition, DLG-1 was usually visibly continuous between the anterior valve and int1, which may explain why their barrier function remained intact. This result suggests that a secondary mechanism may drive later septate-like junction formation independent of PAR-6 and PKC-3. A candidate parallel pathway is the Crumbs polarity complex, as the transmembrane domain protein Crumbs is known to be a critical polarity regulator in many epithelia (Charrier et al., 2015; Tepass et al., 1990). Three Crumbs homologs have been identified in *C. elegans*, and simultaneous removal of all three does not cause obvious polarity defects (Waaijers et al., 2015), but their role could normally be masked by the activity of the PAR complex.

#### Lumen formation

Lumenogenesis requires correct apicobasal polarity, as apically-directed vesicles add membrane, apical proteins, and channel and pump proteins to expand the apical surface and build the luminal space (Datta et al., 2011; Shafaq-Zadah et al., 2020). Yet without PAR-6, larval intestines developed luminal regions surrounded by apical proteins, suggesting that the ability of the apical domain to direct apical trafficking and lumenogenesis was intact. The edematous nature of much of the lumen that formed, particularly in PAR-6^gut(−)^ and PKC-3^gut(−)^ larvae, may indicate an inability to regulate luminal expansion. However, luminal expansion is usually associated with increased apical polarity proteins, such as overexpressed Crumbs, or decreased basolateral proteins (Datta et al., 2011), the opposite of our depletion cases. Therefore, we favor a model in which the edematous swellings are the byproduct of normal lumenogenesis at the apical surface but with no exit route for the luminal fluid, leading to swelling due to trapped fluid accumulation.

An important caveat to our interpretations is that tissue-specific degradation is not equivalent to a null allele. While we see robust degradation of all three proteins (Supp. Fig. 2), we cannot exclude the possibility that low levels of undegraded protein are sufficient to rescue lumenogenesis and barrier function. However, the level of PAR-6 and PKC-3 degradation we achieve is clearly sufficient to cause fully penetrant luminal gaps and larval arrest. Thus, we consider the likeliest explanation to be that PAR-6, PKC-3, and CDC-42 are not required in the developing intestine to build a lumen *per se* or to maintain barrier function. Rather, these proteins maintain apical and junctional domain continuity and thus continuous lumenogenesis, resulting in a functional intestinal tube.

### PAR-6 and PKC-3 play a more substantial role than CDC-42 in maintaining apical continuity

In many tissues, CDC-42 is required for PAR complex localization and to activate PKC-3 kinase activity (Campanale et al., 2017; Pichaud et al., 2019), but CDC-42 is not required to establish apical polarity or to recruit apical PAR-6 in the intestine (Zilberman et al., 2017). Our results revealed two surprising findings about CDC-42. First, by depleting CDC-42 specifically in the intestine, we bypassed the embryonic lethality caused by skin-based enclosure defects (Zilberman et al., 2017), and we were able to identify a later role for CDC-42 in promoting intestinal apical continuity after apical polarity is established. Second, while CDC-42^gut(−)^, PAR-6^gut(−)^ and PKC-3^gut(−)^ embryos all had similar embryonic gap defects, we found that the number of intestinal gaps defects increased only in PAR-6^gut(−)^ and PKC-3^gut(−)^ larvae, and not in CDC-42^gut(−)^ larvae. This result indicates that PAR-6 and PKC-3 continue to be required to maintain apical continuity as the embryonic intestine further elongates, whereas CDC-42 may only be required early in intestinal morphogenesis. One possible explanation for the difference in larval phenotypic severity could be that CDC-42 is specifically important for activating PKC-3 kinase activity, but that PAR-6 and PKC-3 also provide kinase-independent activity, as has been observed in the *Drosophila* embryonic ectoderm (Kim et al., 2009). Consistent with this possibility, the PKC-3 kinase mutant G336N partially rescued PKC-3^gut(−)^, resulting in a phenotype similar to CDC-42^gut(−)^ larvae. As PAR-6 and PKC-3 both have several protein-binding domains (Pickett et al., 2019), the kinase-independent activity could be to recruit other polarity factors to the apical domain later in intestinal development.

Apicobasal polarity regulators are highly conserved, yet differentially required across organisms and tissues (Pickett et al., 2019; Wen and Zhang, 2018). Similarly, the same gene can have oncogenic or tumor suppressive functions in different tissues; for example, *PARD3*/Par3 overexpression is associated with renal cancers, but *PARD3* downregulation or deletion is associated with breast, glioblastoma, lung, and other cancers (Halaoui and McCaffrey, 2015). Detailed studies of diverse epithelia will continue to be important for parsing apart the roles of critical polarity proteins during different stages of organ development and homeostasis, to deepen our understanding about the many ways in which these highly conserved regulators function in development and in disease.

## MATERIALS AND METHODS

### *C*. *elegans* strains and maintenance

Nematodes were maintained at 20°C and cultured and manipulated as previously described (Sulston and Brenner, 1974), unless otherwise stated. Experiments were performed using embryos and larvae collected from 1- or 2-day-old adults. The strains used in this study are listed in Supplemental Table 3.

### New alleles and transgenes

#### CRISPR editing

The self-excision cassette (SEC) method was used to generate CRISPR alleles (Dickinson et al., 2015). GFP or ZF::GFP alleles also contain 3×FLAG and TagRFP-T alleles also contain 3×Myc. The plasmid pDD162 used to deliver Cas9 and each sgRNA was modified (Q5 Site-Directed Mutagenesis Kit, NEB) to insert the appropriate sgRNA guide sequence for each CRISPR edit. The repair template was generated by PCR-amplifying appropriate homology arm sequences for an N-terminal, C-terminal, or internal fluorophore insertions (Phusion High-Fidelity DNA polymerase, Thermo Scientific) and cloned into an SEC backbone plasmid (NEBuilder HiFi DNA Assembly Master Mix, NEB). The modified Cas9/sgRNA plasmid, repair template, and pBS were injected at 50 ng/μL each into N2 or *zif-1(gk117)* mutant 1-day-old adult worms. Injected worms were recovered and treated according to published protocols to isolate independent CRISPR edit events. New CRISPR alleles were backcrossed at least twice before being used for subsequent experiments. The internal in-frame placement of the fluorophore in PAR-3 tags all isoforms without causing an obvious phenotype. PAR-6 was tagged at the C-terminus with ZF::GFP and ZF::GFP_11_. Both *par-6* alleles resulted in full penetrant larval arrest when degraded in the intestine (Fig. 6A). sgRNA and homology arm sequences, plasmids, and primers used in constructing new CRISPR alleles are listed in Supplemental Table 4.

#### Integrated and extrachromosomal arrays

*wowIs3[ifb-2*p::*zif-1*] (IV or V) was derived by spontaneous integration of the extrachromosomal array *wowEx34* (Sallee et al., 2018). *wowIs28[elt-2*p::*zif-1*] (II) was derived by spontaneous integration of an extrachromosomal array carrying SA109 (*elt-2p*::*zif-1* at 50ng/μL (Armenti et al., 2014)), pJF248 (*end-1p*::*histone*::*mCherry* at 50ng/μL), pCFJ90 (*myo-2p*::*mCherry* at 2.5ng/μL (Frøkjær-Jensen et al., 2008)), and pBS (47.5ng/μL). The YFP::ACT-5 transgene *opIs310* was genetically linked to *wowIs3*, so *wowIs28[elt-2*p::*zif-1]* (II) was used instead for intestine-specific ZIF-1 expression.

Extrachromosomal arrays were generated to provide intestine-specific expression of wild-type or mutant PKC-3. The plasmid pMS252 containing *elt-2*p::*bfp::pkc-3::unc-54* 3’UTR was generated by PCR-amplifying *pkc-3* from N2 wild-type genomic DNA and the *unc-54* 3’UTR sequence from pSA109 with Phusion DNA polymerase, and using the NEBuilder Assembly Mix, PCR products were cloned into a plasmid carrying *elt-2*p::*bfp* (pMP27) digested with AfeI and XmaI. Mutations were introduced by Q5 mutagenesis to make pMS259 carrying a PKC-3(G336N) mutation (Forward primer: ATTCGTTCCTaatGGTGATCTGATG; Reverse primer: TCGATGACAAAGAACAGG) and to make pMS260 carrying a deletion of the PKC-3 N-terminus carrying the PB1 domain (Forward primer: AAACCAGAGCTGCCCGGG; Reverse primer: AGCGCTGTTGAGCTTGTGTC). For the PKC-3 rescue assay, *end-1*p::*bfp*::*pkc-3* gDNA plasmids were each injected at 10 ng/μL into JLF148 young adult hermaphrodites along with pBS (115 ng/μL) as carrier DNA and *unc-122*p::GFP (25 ng/μL) to mark the coelomocytes, and independent lines were isolated and crossed into JLF491 to test for function.

### Obtaining gut(−) depletion embryos and larvae

All PAR-6, PKC-3, and CDC-42 intestine-specific depletion strains were maintained with the ZF::GFP allele balanced by the pharyngeal GFP-marked *hT2* (PAR-6) or *mIn1* (PKC-3, CDC-42). To obtain “gut(−)” embryos, hermaphrodite L3 and L4 larvae lacking the balancer were transferred to a fresh plate, and their embryos were scored for defects so that both maternal and zygotic supplies of each protein was ZF::GFP-tagged and susceptible to degradation. Depletion was verified by examining the loss of GFP fluorescence at the intestinal midline (Supp. Fig. 2). While this strategy does not generate a null situation, this system robustly depletes ZF-tagged proteins (Magescas et al.; Sallee et al., 2018; Sanchez et al.), and we see that again here (Supp. Fig. 2), with only occasional and extremely weak midline GFP signal being detectable. This occasional weak signal was only observed when the *wowIs3* transgene began to be silenced over time in a few strains, and could be identified by the visible loss of pharyngeal mCherry expression, a co-injection marker of the transgene. Also, rare “escaper” larvae did not arrest at the L1 stage when silencing occurred.

### Microscopy

Embryos were raised at 20°C and either dissected from gravid hermaphrodites after a 5-hour incubation in M9 for time courses and star cell divisions, or collected from plates containing 1 day old gravid adults. Embryos were mounted on a pad made of 3% agarose dissolved in M9 and imaged using a Nikon Ti-E inverted microscope (Nikon Instruments, Melville, NY) with a 60× Oil Plan Apochromat (NA = 1.4) objective controlled by NIS Elements software (Nikon). Images were acquired with an Andor Ixon Ultra back thinned EM-CCD camera using 488 nm or 561 nm imaging lasers and a Yokogawa X1 confocal spinning disk head equipped with a 1.5× magnifying lens. Images were taken at a z-sampling rate of 0.5 μm. Live time-lapse imaging was done with 4 minute time steps for star cell divisions (Fig. 2) and with 10 minute steps for elongation (Fig. 3).

For fluorescence imaging in larvae, young adult hermaphrodites were allowed to lay eggs overnight, and larvae were picked into a drop of 2mM levamisole on a 3% agarose pad to minimize movement, and imaged on a Nikon Ni-E compound microscope with an Andor Zyla sCMOS camera with a 60X Oil Plan Apochromat (NA = 1.4) objective and NIS Elements software. For imaging larvae for the “Smurf” feeding assay, a Vankey Cellphone Telescope Adapter Mount (Amazon) was used with an Apple iPhone 7 camera and the compound microscope 60X objective. Brightness and hue were adjusted for each image with Adobe Photoshop v21.0.1.

Percent laser power and exposure time were the same for all genotypes imaged within each experiment. Images were processed in NIS Elements and the Fiji distribution of ImageJ (“Fiji”) (Schindelin et al., 2012).

### Image quantification

Quantitative analysis of midline fluorescence signal intensity of MTOC markers TBG-1::mCherry and PTRN-1::GFP in Fig. 3 was performed as follows. For each genotype, embryos were imaged with a 400ms exposure and 80% 488 laser power for PTRN-1::GFP, and a 400ms exposure and 100% 561 laser power for TBG-1::mCherry. Midline signal fluorescence intensity was measured using Fiji by drawing a 4 pixel-wide segmented line on a maximum projection of the z-slices that capture the intestinal midline signal, carefully avoiding the pharyngeal and rectal valve signal coming from just outside the intestine. Intestinal background signal fluorescence intensity was measured by drawing a line of variable length (minimum 10 microns) in the cytoplasm of the intestine, avoiding the midline, the germ cells, and the nuclei. Each measurement was normalized as follows: (measurement – average intestinal background intensity)/(average intestinal background intensity). We reasoned that midline signal intensity in a gap should be in the same range as intestinal background signal intensity, so we used intestinal background variability to define a “background-level” measurement. 97.5% of normalized intestinal background measurements were less than two standard deviations above 0, so we used this cutoff (0 + two standard deviations) to distinguish between a “background-level” and apical enrichment-level measurement at the midline. A normalized midline measurement that was background-level for both TBG-1 and PTRN-1 was considered a background-level measurement in our analysis. We defined a “gap” in the midline as a region of the midline with at least three consecutive background-level measurements. This is likely a conservative measurement of gaps, as some gaps were visible to the eye that were not detected by this analysis (compare Fig. 3F to 4E). We did not distinguish between star cell and non-star cell gaps in this analysis.

For time-lapse imaging of elongation (Fig. 3H), we performed the above analysis at two time points. The first timepoint was the first image frame after the anterior cell divisions had completed, and the second time point was 50 minutes later at approximately 1.5-fold stage.

### L1 arrest assay

To assess larval viability, 10-20 one-day-old gravid hermaphrodites were picked to an NGM plate and allowed to lay eggs for 2-4 hours at 20°C, with three trials per genotype. Adults were removed, and larvae that grew older than the L1 larval stage were counted and removed over the following three days before counting the number of remaining L1 larvae, which usually appeared unhealthy or dead by day three.

For the PKC-3 rescue assay strains, gravid hermaphrodites were allowed to lay eggs for 12 hours to increase the number of array-positive progeny to analyze. Two independently derived transgenic lines were scored to assess the rescuing capacity of wild-type PKC-3 and PKC-3(kinase mutant), and one line for PKC-3(PAR-6-binding mutant) were scored.

### Smurf feeding assay

One-day-old gravid hermaphrodites were allowed to lay eggs overnight, and hatched larvae were picked into a 30 μL drop of standard overnight OP50 bacterial culture. 10 uL of a 20% solution of blue food coloring (FD&C Blue #1 Powder, Brand: FLAVORSandCOLOR.com, Amazon) in water was added at a final concentration of 5%, and worms were incubated in a humid box for three hours before imaging. To collect the larvae, the dye-OP50 solution was transferred to and spread on an NGM plate to allow the larvae to crawl out of the dark blue solution. Larvae were mounted in 2mM levamisole on a 3% agarose pad and imaged as described above. Occasionally, larvae were damaged in the transfer process, so larvae that appeared desiccated or damaged were omitted from the analysis. For the PKC-3 rescue assay strains, mosaicism of the array could be detected because transgenically supplied PKC-3 was BFP-tagged and thus visible when it was present or absent. We could not interpret PKC-3 localization, as even wild-type PKC-3 appeared cytoplasmic, likely due to its overexpression. However, the presence of BFP allowed us to exclude from our analysis any larvae in which mosaicism appeared to correlate with intestinal defects.

### Statistical analysis

Statistical analyses were performed in Excel and RStudio. When comparing multiple genotypes (Figures 2D, 3F, 4E, 7F), we used ANOVA to confirm there were statistically significant differences between groups (p < 10-3), followed by a post-hoc Tukey’s test for pairwise comparisons of the genotypes to determine p-values in the figures. When comparing two genotypes, either a Fisher’s exact test or a two-tailed t-test was used.

## ACKNOWLEDGEMENTS

We thank Jeremy Nance, Dan Dickinson, Ken Kemphues, Victor Naturale, and Lauren Cote for strains and plasmids, and members of the Feldman lab for helpful discussions about the project and manuscript. Some strains were provided by the CGC, which is funded by NIH Office of Research Infrastructure Programs (P40 OD010440). This work was funded by 1K99GM13548901 to M.D.S., F32 GM129900-01 to M.A.P., and an NIH New Innovator Award DP2GM119136-01 and R01GM133950 awarded to J.L.F.

