## Supplemental Material for "Apical PAR complex proteins protect against epithelial assaults to create a continuous and functional intestinal lumen"

**SUPPLEMENTARY MATERIAL**

**Supplementary Figures:**

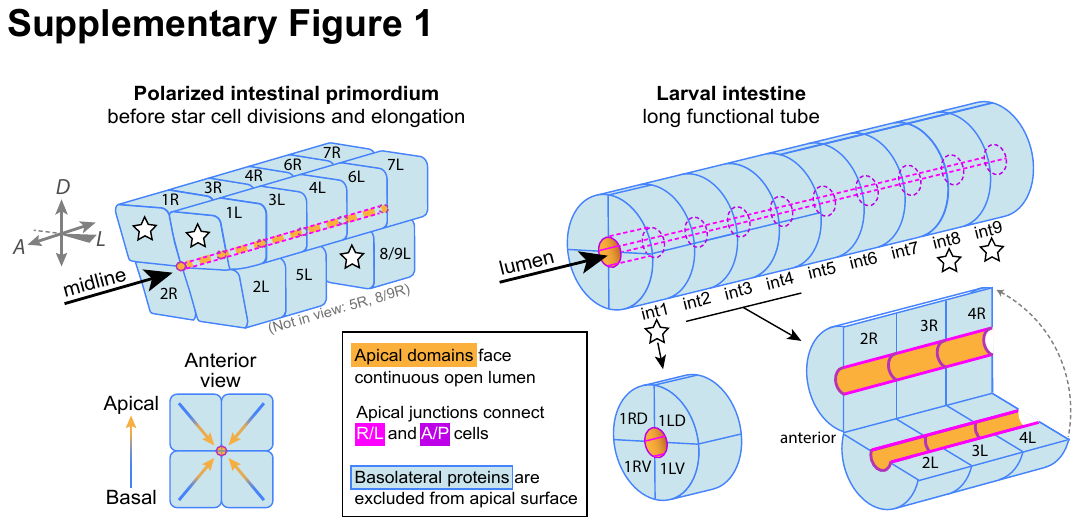

**Supplementary Figure 1.** *C. elegans intestinal development*

*Left*: A 3D representation of the intestinal primordium prior to star cell divisions and elongation, with an end-on anterior view showing the apical domains (gold) of intestinal cells facing the central midline, separated from the basolateral domains (blue) by junctions (magenta). One of the four star cells (8/9R) and one of the twelve non-star cells (5R) are not in view. Numbers indicate which int ring each cell or its descendants will form (eg., 3R and 4R will become part of int3 and int4, respectively). Different shades of magenta are used to highlight different cell-cell interactions. Bright magenta indicates junctions between left and right neighbors, which will together build an int ring; dark magenta indicates junctions between anterior and posterior neighbors, which will join adjacent int rings. *Right*: A 3D representation of the L1 larval intestine. All int rings contain two cells except for the four-celled int1. Apical domains of all cells face the central lumen, and junctions connect left/right pairs of cells within int rings (bright magenta) and also anterior and posterior cells of adjacent int rings (darker magenta), forming a ladder-like pattern.

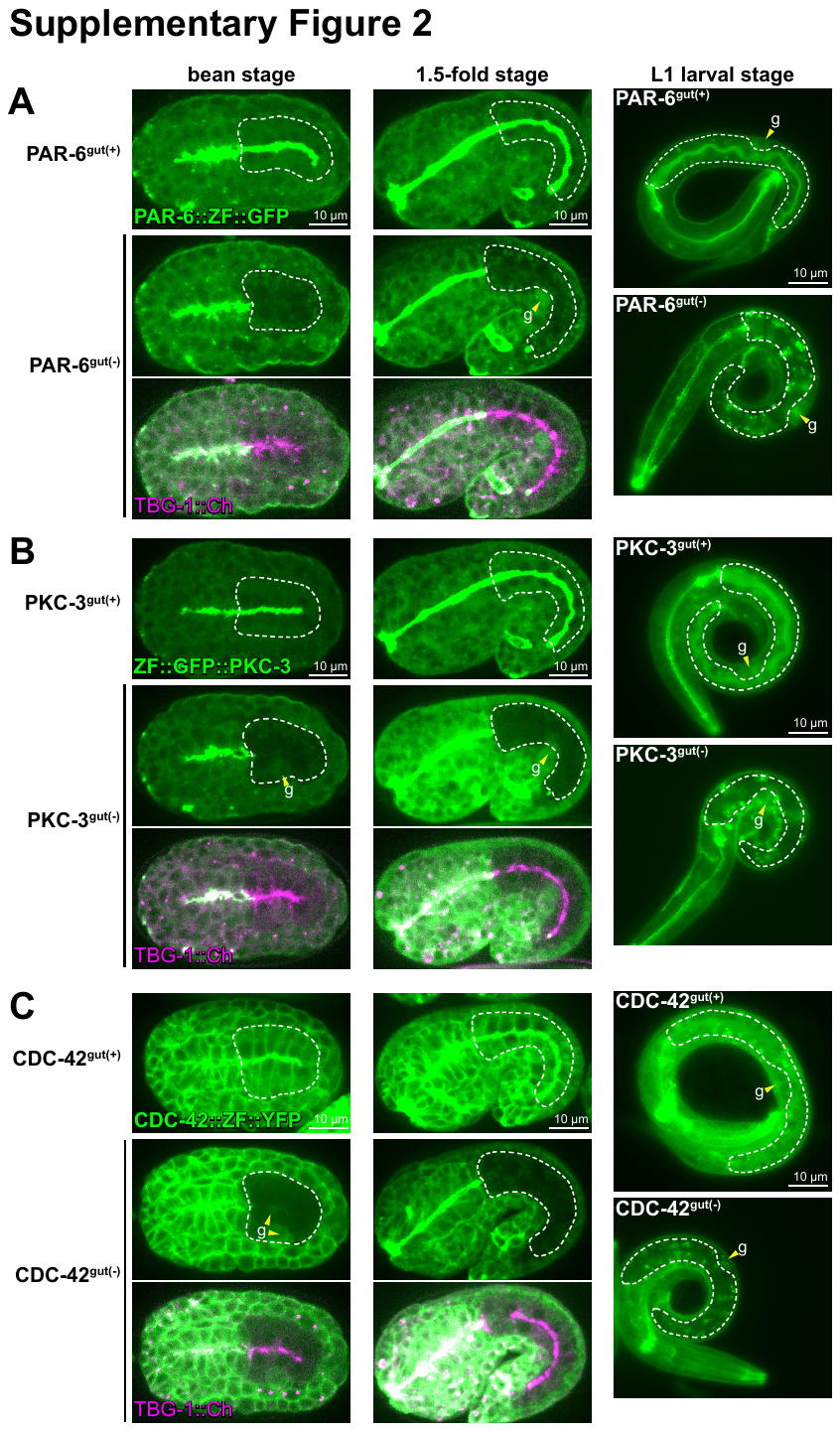

**Supplementary Figure 2.** *ZIF-1/ZF-mediated protein degradation*

Dorsolateral spinning-disk confocal images of bean- and 1.5-fold-stage embryos, and widefield fluorescent images of L1 larvae of the indicated genotypes. “gut(+)” indicates the absence of ZIF-1 and thus that the ZF-tagged protein was not degraded, and “gut(-)” indicates the presence of intestine-specific expression of and degradation by ZIF-1. TBG-1::mCherry was used to mark the intestinal midline in gut(-) embryos. Dashed white lines outline intestines. Yellow arrowheads and “g” indicate germ cells in embryos and the gonad in larvae. Robust depletion by bean stage: PAR-6^gut(-)^: n = 14/15, PKC-3^gut(-)^: n = 25/25, CDC-42^gut(-)^: n = 15/15. In larval intestines, bright green puncta are birefringent “gut granules” and not fluorescent signal from the CRISPR alleles. Scale bars = 10 μm.

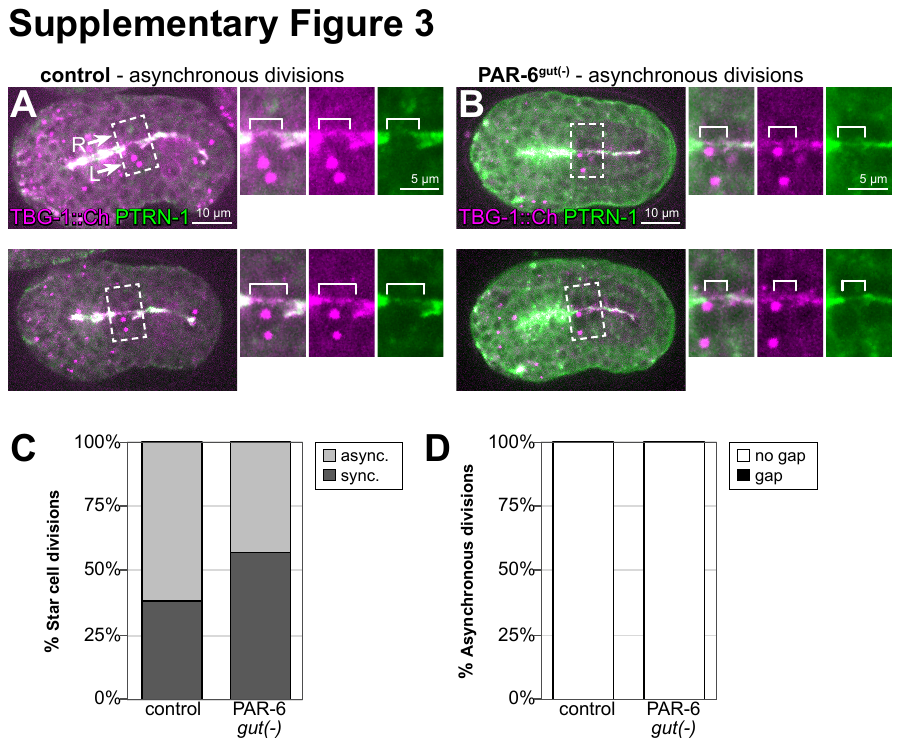

**Supplementary Figure 3.** *Asynchronous star cell divisions in control and PAR-6 embryos do not cause gaps in the apical MTOC.*

Dorsal view confocal images of bean stage embryos in which the star cell divisions were asynchronous. The left star cell (white arrow, “L”) was dividing and the right star cell (white arrow, “R”) was not. PTRN-1::GFP and TBG-1::mCherry mark the apical MTOC. TBG-1 also marks active centrosomes during mitosis. Maximum intensity Z-projections capture the active centrosomes, but not always the entire midline. 2X magnified images of boxed region highlighting midline gaps are shown at right (merged channels, TBG-1::mCherry only, and PTRN-1::GFP only). White brackets indicate midline region of the non-dividing right cell. (A) Two representative control embryos. (B) Two representative PAR-6^gut(-)^ embryos. (C) Graph showing the percent of star cell divisions that are asynchronous versus synchronous in control (n = 13) and PAR-6^gut(-)^ (n = 14) embryos. Two-tailed Fisher’s exact test, p = 0.4495. (D) Graph showing the percent of asynchronous star cell divisions in which a gap forms in control (n = 0/8) and PAR-6^gut(-)^ (n = 0/6) embryos.

**Supplementary Tables:**

**Supplementary Table 1. Larval arrest data from three trials.**

| **Description** | **Strain: genotype of larvae scored** | **Trial** | **L1 after 3 days** | **Older than L1** |
| --- | --- | --- | --- | --- |
| E8 zif-1 | JLF204: *ltSi569; zif-1(gk117); wowIs3* | 1 | 0 | 174 |
|  |  | 2 | 2 | 196 |
|  |  | 3 | 0 | 182 |
| E4 zif-1 | JLF724: *ltSi569 par-6(wow31)/tmC27[myo-2p::Venus]; wowIs28; zif-1(gk117); opIs310* | 1 | 3 | 227 |
|  |  | 2 | 8 | 190 |
|  |  | 3 | 7 | 213 |
| PAR-6::ZG | JLF212: *par-6(wow31); zif-1(gk117)* | 1 | 3 | 185 |
|  |  | 2 | 4 | 166 |
|  |  | 3 | 1 | 168 |
| PAR-6::ZG11 | JLF758: *par-6(wow119); zif-1(gk117)* | 1 | 0 | 195 |
|  |  | 2 | 1 | 183 |
|  |  | 3 | 0 | 187 |
| ZG::PKC-3 | JLF480: *pkc-3(wow85); zif-1(gk117)* | 1 | 2 | 188 |
|  |  | 2 | 1 | 206 |
|  |  | 3 | 1 | 164 |
| ZY::CDC-42 | JLF882: *cdc-42(xn65); zif-1(gk117)* | 1 | 0 | 474 |
|  |  | 2 | 0 | 541 |
|  |  | 3 | 0 | 560 |
| PAR-6::ZG E8 gut(-) | JLF354: *ltSi569 par-6(wow31); zif-1(gk117); wowIs3* from balancer(-) mothers | 1 | 137 | 0 |
|  |  | 2 | 109 | 2 |
|  |  | 3 | 127 | 0 |
| PAR-6::ZG11 E8 gut(-) | JLF726: *par-6(wow119); zif-1(gk117); wowIs3* from balancer(-) mothers | 1 | 167 | 0 |
|  |  | 2 | 107 | 0 |
|  |  | 3 | 141 | 1 |
| PAR-6::ZG E4 gut(-) | JLF724: *ltSi569 par-6(wow31); wowIs28; zif-1(gk117); opIs310* from balancer(-) mothers | 1 | 137 | 0 |
|  |  | 2 | 109 | 2 |
|  |  | 3 | 127 | 0 |
| ZG::PKC-3 E8 gut(-) | JLF491: *pkc-3(wow85); zif-1(gk117); wowIs3* from balancer(-) mothers | 1 | 57 | 3 |
|  |  | 2 | 85 | 3 |
|  |  | 3 | 78 | 3 |
| ZY::CDC-42 E8 gut(-) | JLF883: *ltSi569; cdc-42(xn65); zif-1(gk117); wowIs3* from balancer(-) mothers | 1 | 66 | 111 |
|  |  | 2 | 51 | 84 |
|  |  | 3 | 64 | 86 |
| PKC-3 E8gut(-), PKC-3(+) line 1 | JLF866: *pkc-3(wow85); zif-1(gk117); wowIs3; wowEx143[end-1*p::BFP::PKC-3(+)] "line 1" | 1 | 2 | 54 |
|  |  | 2 | 7 | 55 |
|  |  | 3 | 5 | 34 |
| PKC-3 E8gut(-), PKC-3(+) line 2 | JLF867: *pkc-3(wow85); zif-1(gk117); wowIs3; wowEx144[end-1*p::BFP::PKC-3(+)] "line 2" | 1 | 4 | 117 |
|  |  | 2 | 3 | 104 |
|  |  | 3 | 4 | 101 |
| PKC-3 E8gut(-), PKC-3(G336N) line 1 | JLF868: *pkc-3(wow85); zif-1(gk117); wowIs3; wowEx146[end-1*p::BFP::PKC-3(G336N)] "line 1" from balancer(-) mothers | 1 | 50 | 50 |
|  |  | 2 | 56 | 45 |
|  |  | 3 | 50 | 43 |
| PKC-3 E8gut(-), PKC-3(G336N) line 2 | JLF869: *pkc-3(wow85); zif-1(gk117); wowIs3; wowEx147[end-1*p::BFP::PKC-3(G336N)] "line 2" from balancer(-) mothers | 1 | 37 | 55 |
|  |  | 2 | 51 | 29 |
|  |  | 3 | 55 | 44 |
| PKC-3 E8gut(-), PKC-3(ΔPB) line 1 | JLF870: *pkc-3(wow85); zif-1(gk117); wowIs3; wowEx150[end-1*p::BFP::PKC-3(ΔPB)] "line 1" from balancer(-) mothers | 1 | 113 | 4 |
|  |  | 2 | 97 | 3 |
|  |  | 3 | 72 | 2 |

**Supplementary Table 2. “Smurf” feeding assay data from three trials.**

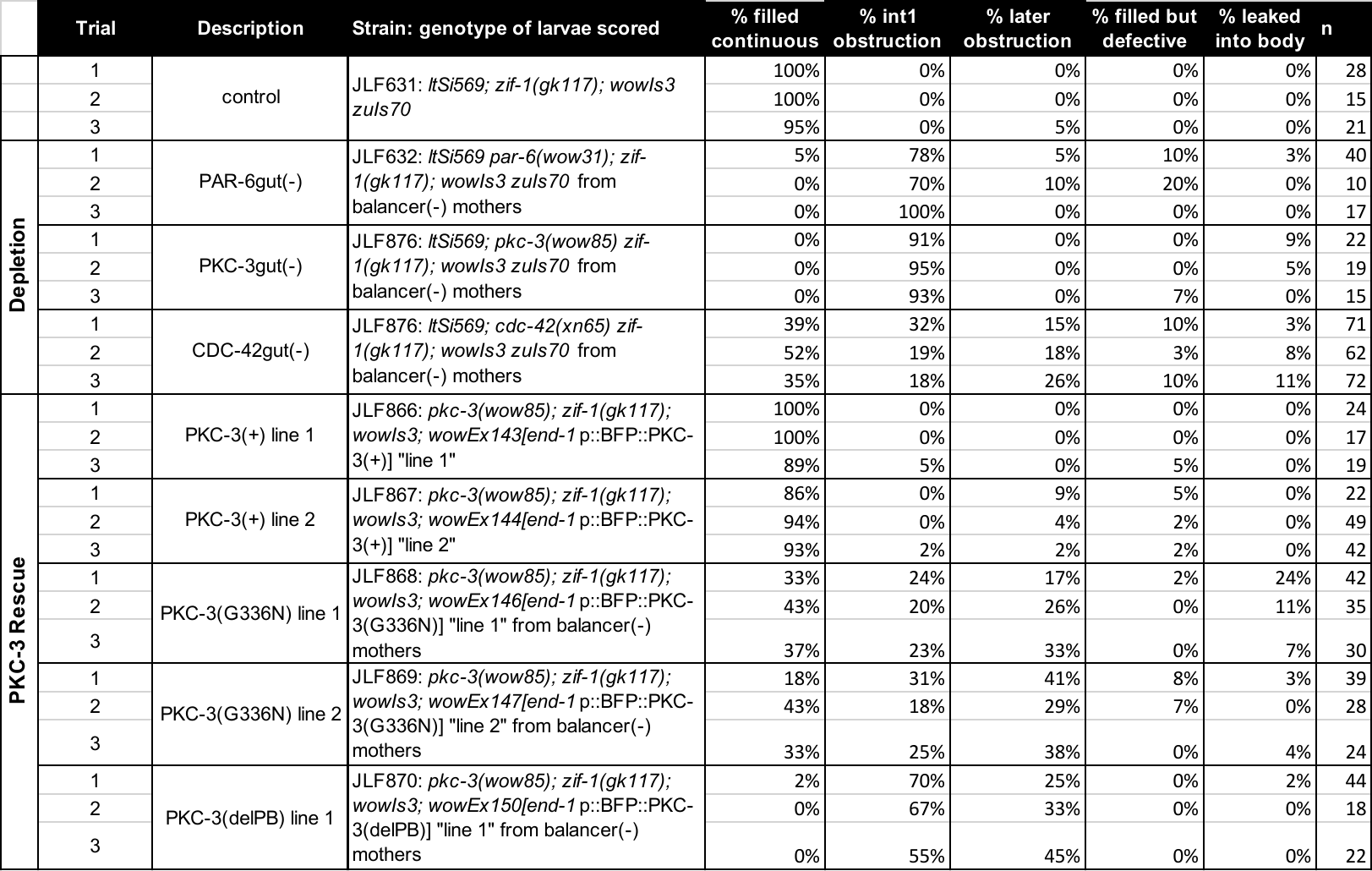

**Supplementary Table 3. Strains used in this study**

| *Figure* | *Strain* | *Genotype* | *Allele source* |
| --- | --- | --- | --- |
| 1B, 1C, 4A, 6C, 6G, 7A | JLF631 | *ltSi569*[TBG-1::mCherry]; *zif-1(gk117)*; *wowIs3*[*ifb-2*p::*zif-1*, *myo-2*p::mCherry] *zuIs70*[*end-1*p::GFP::CAAX] | (Asan et al., 2016; Wang et al., 2015) |
| 1E | JLF37 | *zuIs278*[*pie-*1pmCherry::TBA-1]; *gip-1*(*wow5*[ZF::GFP::GIP-1]) *zif-1(gk117)* | (Sallee et al., 2018) |
| 1E | JLF83 | *ltSi569*; *ptrn-1(wow4*[PTRN-1::GFP]) | (Sallee et al., 2018) |
| 1E | JLF153 | *ltSi569*; *zif-1(gk117)*; *zyg-9*(*wow13*[ZYG-9::ZF::GFP]) | (Sallee et al., 2018) |
| 1E | JJ2376 | *ddIs6*[*pie-1*p::GFP::TBG-1]; *tjIs222*[*pie-1*p::mCherry::AIR-1] |  |
| 1E | JLF152 | *ltSi569*; *zif-1(gk117)*; *noca-1*(*wow11*[NOCA-1::ZF::GFP]) | (Sallee et al., 2018) |
| 1E | JLF729 | *ltSi569* *vab-10*(*wow80*[VAB-10B::ZF::GFP]); *zif-1(gk117)* | (Sanchez et al.) |
| 1F | JLF878 | *ltSi569*; *zif-1(gk117)*; *wowIs3*; *dlg-1(cp301*[DLG-1::mNeonGreen]) | (Heppert et al., 2018) |
| 1F | JLF150 | *ltSi569*; *unc-119(ed3)*; *xnIs96*[HMR-1::GFP] | (Achilleos et al., 2010) |
| 1G | JLF442 | *zuIs278*; *opIs310*[*ced-1*p::YFP::ACT-5] | (Neukomm et al., 2011) |
| 1H | JLF148 | *ltSi569* *par-6(it319*[PAR-6::GFP]); *unc-119(ed3* or *+)* | *CGC* |
| 1H | JLF149 | *ltSi569*; *pkc-3(it309*[GFP::PKC-3]); *unc-119(ed3)* or *unc-119(+)* | *CGC* |
| 1H | JLF147 | *ltSi569*; *par-3(it298*[PAR-3::GFP]); *unc-119(ed3*) or *unc-119(+)* | *CGC* |
| 2A, 2F, 3A, 3D, S3A | JLF440 | *ltSi569*; *zif-1(gk117)*; *wowIs3*; *ptrn-1(wow4)* |  |
| 2B, 2G, 3B, 3E, S3B | JLF445 | *ltSi569* *par-6(wow31*[PAR-6::ZF::GFP])/*hT2[qIs48]*; *zif-1(gk117)/hT2*; *wowIs3*; *ptrn-1(wow4*) |  |
| 2C, 3C | JLF490 | *ltSi569*; *pkc-3(wow85*[ZF::GFP::PKC-3])/*mIn1*[*mIs14*]; *zif-1(gk117);* *wowIs3; ptrn-1(wow4)* |  |
| 4B, 6D, 6G, 7B | JLF632 | *ltSi569* *par-6(wow31)/hT2[qIs48]*; *zif-1(gk117)/hT2*; *wowIs3* *zuIs70* |  |
| 4C, 6E, 6G, 7C | JLF876 | *ltSi569;* *pkc-3(wow85)/mIn1[mIs14]*; *zif-1(gk117)*; *wowIs3* *zuIs70* |  |
| 4D, 6F, 6F', 6G, 7D | JLF877 | *ltSi569;* *cdc-42(xn65[ZF::YFP::CDC-42])/mIn1[mIs14]*; *zif-1(gk117)*; *wowIs3* *zuIs70* | (Zilberman et al., 2017) |
| 5A, 6A, 8D, 8E | JLF724 | *ltSi569 par-6(wow31)*/*tmC27*[*unc-75*(*tmIs1239*[*myo-2*p::Venus])]; *wowIs28*; *zif-1(gk117)*; *opIs310* | CGC |
| 5C | JLF784 | *par-3(wow121*[PAR-3::tagRFP]) *zif-1(gk117); wowIs3; wow4* |  |
| 5C | JLF785 | *par-6(wow119*[PAR-6::ZF::GFP11])/*hT2[qIs48]*; *zif-1(gk117)/hT2; wowIs3; wow4* |  |
| 5D | JLF719 | *hmr-1(cp21*[HMR-1::GFP]) *ltSi569*; *zif-1(gk117)*; *wowIs3* | (Marston et al., 2016) |
| 5D | JLF720 | *hmr-1(cp21)* *ltSi569 par-6*(*wow31)*/*hT2[qIs48]*; *zif-1(gk117)/hT2; wowIs3* |  |
| 5E, 8H | JLF721 | *ltSi569 par-6(wow31)/hT2[qIs48]; zif-1(gk117)/hT2; wowIs3; dlg-1(cp301)* |  |
| 5G | JLF879 | LET-413::GFP *par-3(wow121*[PAR-3::tagRFP]) *zif-1(gk117)*; *wowIs3* | (Legouis et al., 2000) |
| 5G | JLF817 | *par-6(wow119)/hT2[qIs48]*; LET-413::GFP *par-3(wow121*) *zif-1(gk117)/hT2; wowIs3* |  |
| 5H, 8J | JLF880 | *ltSi569; itIs256*[LGL-1::GFP]; *zif-1(gk117); wowIs3* | (Beatty et al., 2010) |
| 5H, 8K | JLF881 | *ltSi569 par-6(wow31)/hT2[qIs48]; itIs256; zif-1(gk117)/hT2; wowIs3* |  |
| 6A | JLF204 | *ltSi569; zif-1(gk117); wowIs3* |  |
| 6A, S2A | JLF212 | *par-6(wow31); zif-1(gk117)* |  |
| 6A | JLF758 | *par-6(wow119); zif-1(gk117)* |  |
| 6A, S2B | JLF480 | *pkc-3(wow85); zif-1(gk117)* |  |
| 6A, S2C | JLF882 | *cdc-42(xn65); zif-1(gk117)* |  |
| 6A, S2A | JLF354 | *ltSi569 par-6(wow31)/hT2[qIs48]; zif-1(gk117)/hT2; wowIs3* |  |
| 6A | JLF726 | *par-6(wow119)/hT2[qIs48]; zif-1(gk117)/hT2; wowIs3* |  |
| 6A | JLF491 | *pkc-3(wow85)/mIn1[mIs14]; zif-1(gk117); wowIs3* |  |
| 6A, S2C | JLF883 | *ltSi569; cdc-42(xn65)/mIn1[mIs14]; zif-1(gk117); wowIs3* |  |
| 6A, 6G, 6H | JLF866 | *pkc-3(wow85); zif-1(gk117); wowIs3; wowEx143[end-1*p::BFP::PKC-3(+)] "line 1" |  |
| 6A, 6G | JLF867 | *pkc-3(wow85); zif-1(gk117); wowIs3; wowEx144[end-1*p::BFP::PKC-3(+)] "line 2" |  |
| 6A, 6G , 6I' | JLF868 | *pkc-3(wow85)/mIn1[mIs14]; zif-1(gk117); wowIs3; wowEx146[end-1*p::BFP::PKC-3(G336N)] "line 1" |  |
| 6A, 6G, 6I | JLF869 | *pkc-3(wow85)/mIn1[mIs14]; zif-1(gk117); wowIs3; wowEx147[end-1*p::BFP::PKC-3(G336N)] "line 2" |  |
| 6A, 6G, 6J | JLF870 | *pkc-3(wow85)/mIn1[mIs14]; zif-1(gk117); wowIs3; wowEx150[end-1*p::BFP::PKC-3(ΔPB)] "line 1" |  |
| 8A | JLF883 | *ltSi569; par-3(wow120*[PAR-3::GFP]) *zif-1(gk117); wowIs3* |  |
| 8B | JLF884 | *ltSi569 par-6(wow31)/hT2[qIs48]; par-3(wow120) zif-1(gk117)/hT2; wowIs3* |  |
| S2B | JLF885 | *ltSi569; pkc-3(wow85)/mIn1[mIs14]; zif-1(gk117); wowIs3* |  |

**Supplementary Table 4. CRISPR allele primers and plasmids**

| *Edit* | **PAR-6::ZF::GFP (C-term)** |
| --- | --- |
| *Allele* | *wow31* |
| *Repair Template* | pMS237 |
| *Repair Template source* | JF250 |
| *5' HA Fwd* | ACGTTGTAAAACGACGGCCAGTCGCCGGCAacgaccacgaaattggctttcg |
| *5' HA Rev* | CATCGATGCTCCTGAGGCTCCCGATGCTCCGTCCTCTCCCgaaTCtGAATCATTTGCGTCGTGCTG |
| *3'Fwd HA* | CGTGATTACAAGGATGACGATGACAAGAGATGAaaaactcttttcagccatttttcctcg |
| *3'Rev HA* | TCACACAGGAAACAGCTATGACCATGTTATtcaaaaatgtccaaatggagcagtgg |
| *sgRNA/Cas9 plasmid 1* | pMS235 |
| *sgRNA guide sequence 1* | GACGCAAATGATTCGGACAG |
| *sgRNA/Cas9 plasmid 2* | pMS236 |
| *sgRNA guide sequence 2* | ACAGCACGACGCAAATGATT |
| *Edit* | **PAR-6::ZF::GFP(11) (C-term)** |
| *Allele* | *wow119* |
| *Repair Template* | pMS255 |
| *Repair Template source* | pLC01 |
| *5' HA Fwd* | ACGTTGTAAAACGACGGCCAGTCGCCGGCAacgaccacgaaattggctttcg |
| *5' HA Rev* | CATCGATGCTCCTGAGGCTCCCGATGCTCCGTCCTCTCCCgaaTCtGAATCATTTGCGTCGTGCTG |
| *3'Fwd HA* | CGTGATTACAAGGATGACGATGACAAGAGATGAaaaactcttttcagccatttttcctcg |
| *3'Rev HA* | TCACACAGGAAACAGCTATGACCATGTTATtcaaaaatgtccaaatggagcagtgg |
| *sgRNA/Cas9 plasmid 1* | pMS235 |
| *sgRNA guide sequence 1* | GACGCAAATGATTCGGACAG |
| *sgRNA/Cas9 plasmid 2* | pMS236 |
| *sgRNA guide sequence 2* | ACAGCACGACGCAAATGATT |
| *Edit* | **ZF::GFP::PKC-3 (N-term)** |
| *Allele* | *wow85* |
| *Repair Template* | pMP41 |
| *Repair Template source* | JF250 |
| *5' HA Fwd* | ttgtaaaacgacggccagtcgccggcagtcttgttcaggccttccag |
| *5' HA Rev* | ACAAAGTCGCGTTTTGTATTCTGTCGGCATtctgaaataaaaattgaactaaaaac |
| *3'Fwd HA* | CGTGATTACAAGGATGACGATGACAAGAGATCGTCTCCGACATCATTAGAAGAGGAC |
| *3'Rev HA* | tcacacaggaaacagctatgaccatgttatTCTGGTTTTCCAACAAAAACG |
| *sgRNA/Cas9 plasmid* | pMP42 |
| *sgRNA guide sequence* | TCGTCTCCGACATCATTAG |
| *Edit* | **PAR-3::GFP (internal)** |
| *Allele* | *wow120* |
| *Repair Template* | pVN36 |
| *Repair Template source* | pDD282 |
| *5' HA Fwd* | acgttgtaaaacgacggccagtcgccggcaTGTCAACCATCAGGCTCACC |
| *5' HA Rev* | TCCAGTGAACAATTCTTCTCCTTTACTGAAAGTGACAAGCGACGTTCC |
| *3'Fwd HA* | CGTGATTACAAGGATGACGATGACAAGAGACCACCGATTCCAGAAAAATCA |
| *3'Rev HA* | tcacacaggaaacagctatgaccatgttatAGTTTCACGTCCGGTCACAG |
| *sgRNA/Cas9 plasmid* | pMP15 |
| *sgRNA guide sequence* | TGATTTTTCTGGAATCGGT |
| *Edit* | **PAR-3::tagRFP (internal)** |
| *Allele* | *wow121* |
| *Repair Template* | pVN39 |
| *Repair Template source* | pDD286 |
| *5' HA Fwd* | cacgacgttgtaaaacgacggccagtcgTGTCAACCATCAGGCTCACC |
| *5' HA Rev* | CTTGATGAGCTCCTCTCCCTTGGAGACGAAAGTGACAAGCGACGTTCC |
| *3'Fwd HA* | GAGCAGAAGTTGATCAGCGAGGAAGACTTGCCACCGATTCCAGAAAAATCA |
| *3'Rev HA* | tcacacaggaaacagctatgaccatgttatAGTTTCACGTCCGGTCACAG |
| *sgRNA/Cas9 plasmid* | pMP15 |
| *sgRNA guide sequence* | TGATTTTTCTGGAATCGGT |
